# Propagation of orientation selectivity in a spiking network model of layered primary visual cortex

**DOI:** 10.1101/425694

**Authors:** Benjamin Merkt, Friedrich Schüßler, Stefan Rotter

## Abstract

Neurons in different layers of sensory cortex generally have different functional properties. But what determines firing rates and tuning properties of neurons in different layers? Orientation selectivity in primary visual cortex (V1) is an interesting case to study these questions. Thalamic projections essentially determine the preferred orientation of neurons that receive direct input. But how is this tuning propagated though layers, and how can selective responses emerge in layers that do not have direct access to the thalamus? Here we combine numerical simulations with mathematical analyses to address this problem. We find that a large-scale network, which just accounts for experimentally measured layer and cell-type specific connection probabilities, yields firing rates and orientation selectivities matching electrophysiological recordings in rodent V1 surprisingly well. Further analysis, however, is complicated by the fact that neuronal responses emerge in a dynamic fashion and cannot be directly inferred from static neuroanatomy, as some connections tend to have unintuitive effects due to recurrent interactions and strong feedback loops. These emergent phenomena can be understood by linearizing and coarse-graining. In fact, we were able to derive a low-dimensional linear dynamical system effectively describing stimulus-driven activity layer by layer. This low-dimensional system explains layer-specific firing rates and orientation tuning by accounting for the different gain factors of the aggregate system. Our theory can also be used to design novel optogenetic stimulation experiments, thus facilitating further exploration of the interplay between connectivity and function.

**Author summary:** Understanding the precise roles of neuronal sub-populations in shaping the activity of networks is a fundamental objective of neuroscience research. In complex neuronal network structures like the neocortex, the relation between the connectome and the algorithm implemented in it is often not self-explaining. To this end, our work makes three important contributions. First, we show that the connectivity extracted by anatomical and physiological experiments in visual cortex suffices to explain important properties of the various sub-populations, including their selectivity to visual stimulation. Second, we introduce a novel system-level approach for the analysis of input-output relations of recurrent networks, which leads to the observed activity patterns. Third, we present a method for the design of future optogenetic experiments that can be used to devise specific stimuli resulting in a predictable change of neuronal activity. In summary, we introduce a novel framework to determine the relevant features of neuronal microcircuit function that can be applied to a wide range of neuronal systems.

## Introduction

Understanding the complex computations performed by neural networks in the nervous system is a central challenge in neuroscience. In recent years, research in rodent sensory systems has provided access to many new facts, but has also raised new theoretical questions. In particular, the role of cortical layers (Niell and Stryker, 2008; Potjans and Diesmann, 2014) and other distinct subpopulations (Durand et al., 2016; Pakan et al., 2016) has received increasing attention. While the anatomical path of sensory input from the thalamus through cortical layers has been particularly well characterized for the visual system, it provides little explanation yet how the information is processed and transformed in transit. Here, we apply computational neuroscience techniques to gain new insight into the various computational steps within a highly recurrent neural network.

From a theoretical perspective, the cortical network constitutes a dynamical system, which operates on different types of sensory input. To study basic properties of such networks, its input has been assumed to be identical for all neurons (Amit and Brunel, 1997; Brunel, 2000). In a sensory system, however, the input varies across neurons, as sensors extract different aspects of the stimulus. The primary visual cortex (V1) represents an interesting example to study such systems (Hansel and van Vreeswijk, 2012; Sadeh et al., 2014). In this case, non-homogeneous input for example carries information about the orientation of moving gratings or light bars presented to the eye of an animal (Hubel and Wiesel, 1959).

A major difference of rodent visual cortex compared to carnivores and primates is the absence of orientation columns and maps (Ohki et al., 2005). Although the “salt-and-pepper” organization of orientation preferences implies that lateral interactions are quite unspecific (Jia et al., 2010; Ko et al., 2011), neurons nevertheless exhibit responses that are strongly tuned with respect to oriented light bars or gratings (Niell and Stryker, 2008; Hofer et al., 2011; Durand et al., 2016). Interestingly, this output tuning is already present at eye opening in juvenile mice, where recurrent connectivity appears to be functionally random (Ko et al., 2013). In recent theoretical studies, this phenomenon could be explained by a strong attenuation of the untuned component of the distributed input, due to the dominance of inhibition in the recurrent network (Hansel and van Vreeswijk, 2012; Pehlevan and Sompolinsky, 2014; Sadeh et al., 2014; Sadeh and Rotter, 2014, 2015). While these previous works identified the mechanism underlying strong output tuning in the primary visual cortex of rodents, they provide no explanation for the different degrees of orientation selectivity in different sub-populations, like cortical layers.

In the present work, we apply computational network modeling techniques to study the emergence of orientation selectivity in a layered cortical network model, with a focus on the differences between layers and neuronal subpopulations. As a starting point, we adopt the anatomically and physiologically founded network model developed by Potjans and Diesmann (2014) and extend it by orientation selective input similar to Sadeh et al. (2014). We model an experiment in which an animal is shown a sinusoidal grating as a visual stimulus. By this approach, we can focus on the influence of synaptic connectivity on neuronal tuning as all other parameters are chosen to be the same over all populations.

Without any adaptation or fine-tuning of parameters, the resulting model matches the distributions of orientation selectivity found in electrophysiological recordings, including the weak tuning of inhibitory neurons as well as L5 pyramidal neurons. We then show that these specific properties cannot be attributed to any specific projection in the circuit.

Rather, it is an emergent network feature of the entire recurrent microcircuit. Furthermore, we suggest a concept for the design of novel optogenetic stimulation experiments. Applying this concept to the V1 cortical microcircuit, we can make specific predictions for the outcome of experiments to manipulate the orientation selectivity of L5 principal cells. The analysis of such experiments can eventually confirm or refute our analysis of the circuit.

## Methods

### A multiplayer model of primary visual cortex

The network model considered throughout our study is, in essence, the same as the recurrent network model developed by Potjans and Diesmann (2014), extended by thalamic input with a weak orientation bias as described in Sadeh et al. (2014). Potjans and Diesmann (2014) created a full-scale neural network model, matching as close as possible the nervous tissue underneath 1 mm^2^ of neocortex. They incorporated a large set of experimental studies, most prominently the ones by Thomson et al. (2002) and Binzegger et al. (2004). They derived a generic connectivity map between eight populations of neurons situated in four layers (L2/3, L4, L5, L6), one excitatory (e) and one inhibitory (i) population in each layer (see Fig 1A for an illustration of the model). The model network consists of close to 80 000 leaky integrate-and-fire neurons, where the respective size of individual subpopulations varies strongly across layers. While the two largest populations (L4e and L2/3e) contain more than 20 000 neurons, L5e comprizes only about 5 000 neurons (Fig 1D). Throughout all layers, inhibitory populations are considerably smaller than the respective excitatory populations, by a factor of four on average. In our work, the leaky integrate-and-fire neurons are all equipped with delta synapses, i.e. with each incoming spike, the membrane potential is instantaneously deflected by a fixed voltage *J*, decaying with the membrane time constant *τ*. The synaptic efficacy *J* varies depending on the source and target population, respectively.

**Figure 1:**
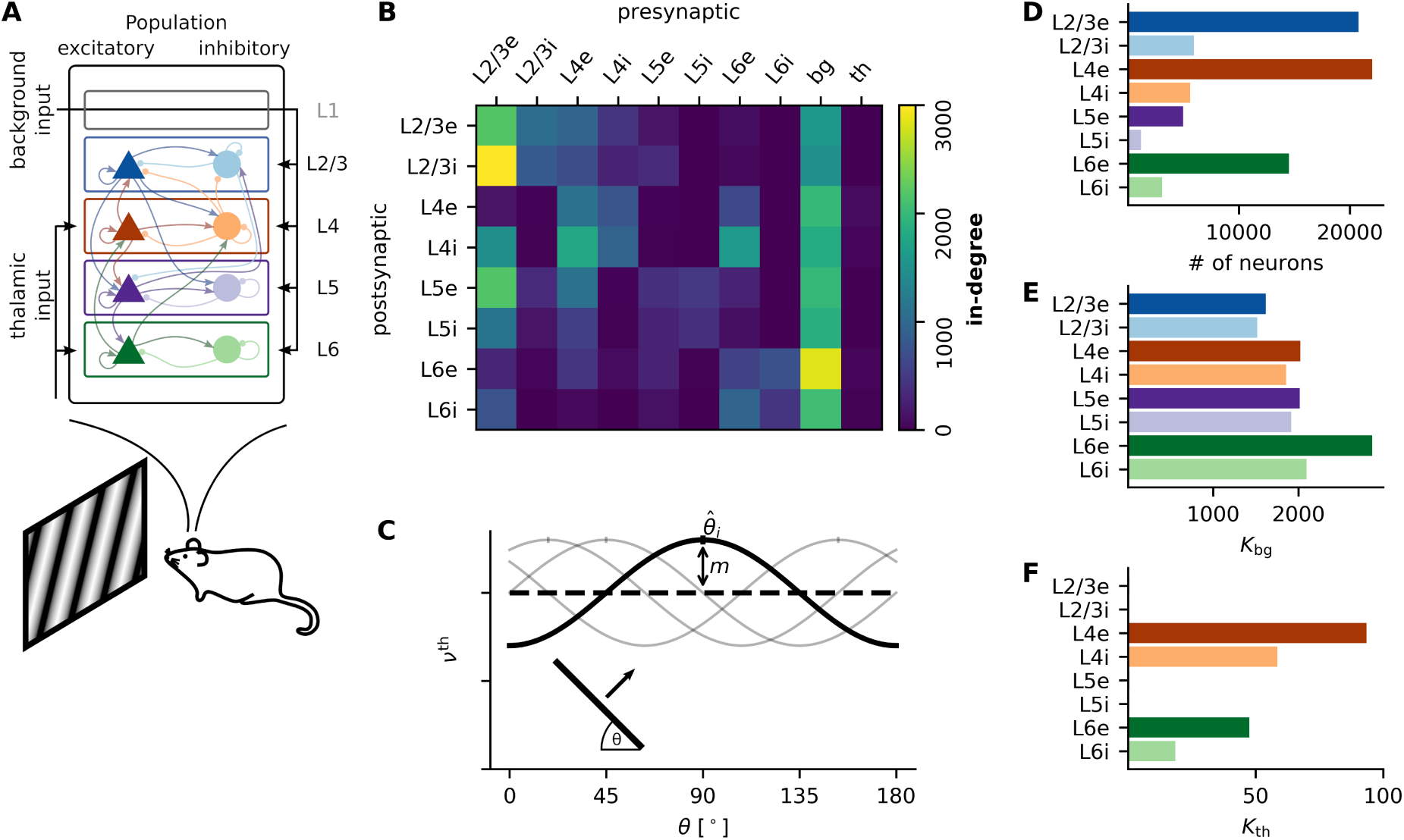
Model parameters. **A** Schematic of the model with eight populations in four layers, thalamic and other background input. Triangles and circles denote excitatory and inhibitory populations, respectively. **B** Total number of synapses each neuron receives (in-degree) from any other neuronal population in the model, including input. **C** Input rate variation over stimulus angles for single neurons in L4 and L6. The angle corresponding to the peak of the curve is the preferred orientation of the thalamic input to that neuron. **D** Total number of neurons in each of the eight subpopulations. **E-F** Input in-degree for each neuron in each of the eight populations. Numbers are identical to the two right-most columns of the matrix shown in B.

Our model has a fixed number of synapses for each projection, i.e. each neuron of the postsynaptic population *P* receives the same number of synaptic inputs *K*_*PT*_ from the presynaptic population *T*. For any postsynaptic neuron in the target population, the presynaptic neurons are drawn randomly from the source population. The numbers *K*_*PT*_, which are the essential parameters of our model, were derived by Potjans and Diesmann (2014) and are graphically represented in Fig 1B.

In addition to recurrent synaptic connections, neurons of the model network receive external input from two different sources (Fig 1A). The first type of input, termed “background input” throughout this paper, is targeting all populations. It represents axons from other cortical and subcortical regions, including gray matter and white matter projections (Fig 1B/E), but excludes those originating from the thalamus. These background inputs are independent of the stimulus. In all simulations, they are modeled as Poisson processes with rate *ν*_*bg*_.

The second type of input is represented by thalamocortical projections that originate from the lateral geniculate nucleus (LGN) (Fig 1B/F). In primary visual cortex, these projections are the main source of information about a stimulus. This is the component where we made an important addition to the original model of Potjans and Diesmann (2014). We replaced the unspecific thalamic input with an input that depends in a specific way on the stimulus. Throughout this paper, the stimuli considered were oriented moving gratings, which are commonly used in visual neuroscience. Similar to Sadeh et al. (2014), each neuron in one of the two populations that receive input (i.e. L4 and L6) is randomly assigned a preferred orientation 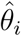 to represent the tuning of the effective compound input from all pre-synaptic thalamic neurons. Specifically, the input to each neuron depends on the orientation *θ* of the stimulus and varies according to

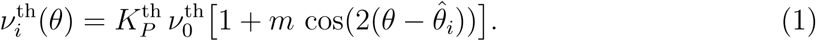

Here, *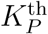* is the in-degree of this projection, and *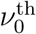* is the mean rate of individual thalamic neurons. We only require the compound input to be tuned and do not make any assumptions about the origin of this tuning. An illustration of the input variation is provided in Fig 1C. Note that orientations vary between 0*°* and 180*°* as we do not consider direction selectivity in this study. Similar to the background input, the thalamic input was conceived as a homogeneous Poisson process.

It is instructive to analyze and compare two different modes of operation. The *stimulation* condition is the situation described above, emphasizing the presentation of a structured visual stimulus (here, a moving oriented grating) to the animal. In contrast, the *spontaneous* condition emulates the presentation of a homogeneous gray screen to the animal. In this case, the intensity of thalamic inputs is the same as the rate of the background inputs. Moreover, the angular modulation *m* is set to zero, removing any information about stimulus orientation from the input.

### Implementations of the model

The results of this study are based on three different adaptations of the network model. Each of them makes additional assumptions about the network dynamics to achieve different levels of abstraction and simplification. At the same time, all renderings are based on the same connectivity parameters and the same single neuron model. In the following, the three models will be briefly described.

#### Model A: spiking neuronal network model

The starting point for our analysis is a *spiking neuronal network*, very similar to the implementation by Potjans and Diesmann (2014). Here, for each of the close to 80 000 neurons, the temporal evolution of the membrane potential *V*_*i*_(*t*) is simulated for each neuron *i* using standard numerical methods. The interaction among neurons in the network is mediated by discrete spike events. Each spike emitted by a pre-synaptic neuron depolarizes or hyperpolarizes the membrane potential of the post-synaptic neuron. The output of the recurrent network is defined in terms of the spike trains of all neurons. Like Potjans and Diesmann (2014), we use the spiking network simulation tool NEST (Gewaltig and Diesmann, 2007) to perform all the necessary numerical simulations.

#### Model B: nonlinear firing rate model

Based on the spiking neural network model, we also developed a neuron-by-neuron *firing rate model*, applying the theory developed by Brunel (2000) to single neurons (Ostojic, 2014; Sadeh and Rotter, 2015). This approximation assumes that neuronal spike trains have Poissonian statistics and that correlations among neurons are insignificant.

The network is characterized by the input-output transfer function *F*_*i*_ of each neuron, leading to the network-level self-consistency equation

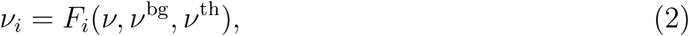

where *i* = 1, *…*, *N* is the neuron index and *ν, ν*^bg^ and *ν*^th^ are the vectors of recurrent neuronal firing rates and input rates, respectively. Throughout this work, variables denoted by th and bg refer to quantities originating from the thalamus or other external inputs, respectively. Consequently, the model results in a high-dimensional system of nonlinear algebraic equations to be solved numerically, yielding the firing rate *ν*_*i*_ of each individual neuron.

#### Model C: linearized network model

Finally, based on Model B, we also derive a *linearized network model* of the microcircuit. Fixing a point of linearization, for any input perturbation Δ*ν*^th^, it predicts the perturbation of the single neuron firing rates Δ*ν*. In the linear model, these can be explicitly calculated by

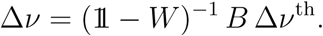

The matrices *W* and *B* are given by appropriate partial derivatives of the transfer function (Eq. 2). As the linearization point, we choose the activity into which the network settles when thalamic neurons are completely silenced. This choice provides the greatest flexibility in the further analysis. Consequently, with this linearization point, we have Δ*ν*^th^ = *ν*^th^ (Eq. 1).

The perturbation Δ*ν*^th^ has different values for the two simulated conditions. In the spontaneous condition, it is given by a homogeneous vector with the compound thalamic baseline rate in all entries. In the stimulated condition, it contains the homogeneous (constant) as well as the non-homogeneous (proportional to *m*) terms in Eq. 1. The latter, therefore, also represents the specific, orientation dependent perturbation for each neuron.

While the spiking network model (Model A) is the most realistic version from a biological perspective, accounting for irregular spike trains and activity fluctuations, it is at the same time computationally expensive and analytically intractable. The firing rate model (Model B) makes additional assumptions about the network activity, like negligible correlations, but it has the advantage of being numerically tractable. Finally, while the linear network model (Model C) ignores non-linear effects by construction, it has the great benefit of allowing the application of a powerful linear algebra toolbox and enabling an explicit solution of the system (Sadeh and Rotter, 2015).

Importantly, considering different implementations of the same model is only justified if their predictions are consistent. Therefore, whenever possible throughout this work, we cross-check the results of the different models.

### Parameter variation

In addition to the standard parameter set adopted from Potjans and Diesmann (2014), we also performed simulations with crucial model parameters varied. Hereby, we were able to evaluate the robustness of our results, and of the conclusions drawn from them. In total, we used six different parameter sets:

- *Standard:* Default parameter set described below.
- *High background:* Background firing rate *ν*^bg^ increased by a factor of 2.
- *Low background:*Background firing rate *ν*^bg^ reduced by a factor of 2.
- *Strong inhibition:* Relative strength of inhibitory synapses increased by a factor of 2.
- *Strong synapses:* Strength of all recurrent synapses increased by a factor of 2.
- *Low connectivity:* Recurrent indegrees reduced by a factor of 2.

## Data analysis

### Measuring orientation selectivity

In order to quantify orientation selectivity, we adopt a measure from circular statistics (Batschelet, 1981; Piscopo et al., 2013). If *ν*_*k*_(*θ*) is the firing rate of neuron *k* for stimulation angle *θ*, the orientation selectivity vector (OSV) is calculated by

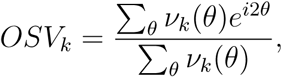

which yields a complex number, equivalent to a vector with two real components. Here, the two sums are running over all stimulation angles *θ*, or a uniform sample of angles in a simulation. The orientation selectivity index (OSI) is then defined as the length of the OSV

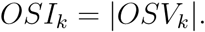

The preferred orientation (PO) of a single neuron can be calculated from the angle of the OSV

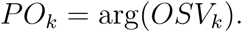

Note that this definition of OSI is identical to

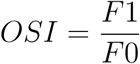

where *F* 0 and *F* 1 are the zeroth and first Fourier components of the tuning curve, giving rise to a more intuitive interpretation of the OSI. Here, *F* 0 accounts for the average firing rate over all stimulation angles, and *F* 1 is to the amplitude of a cosine function accounting for the lowest-order modulation over all angles. A perfect cosine tuning curve as assumed for the input provided by thalamic neurons yields 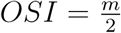, where *m* is the modulation amplitude of the tuning curve (Eq. 1). The OSI as defined here provides a robust measure of tuning strength, in particular for neurons with low firing rates and noisy responses.

An alternative measure for orientation selectivity also used in the literature is based on the neuronal firing rates at the preferred and orthogonal orientation

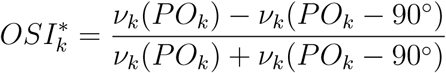

For noisy measurements, in particular for low firing rates, an estimation according to this recipe is problematic. Therefore, we fit a cosine function with offset to the tuning curve of each individual neuron and extract the maximum and minimum firing rates from this fit.

### Input currents

Our model uses a simple integrate-and-fire neuron model with delta-synapses. In this model, the mean current received by neuron *i* through synaptic input from neuron *j* is given by

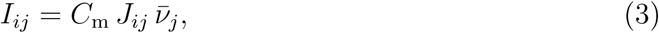

where 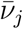 is the mean rate of neuron *j*. In order to quantify the information about the stimulus contained in the input current received from any neuron in the network, we also define a tuning vector (TV) of each synaptic connection. For a given pre-synaptic neuron *j* and post-synaptic neuron *i*, it is defined by

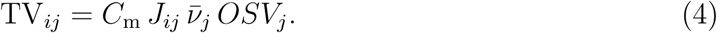

where *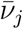* is the firing rate of neuron *j* averaged over all angles. Note that this is identical to the first Fourier component (F1) of the tuning curve of the input current. The single neuron tuning vectors TV_*ij*_ can then be used to quantify the combined tuning information each neuron in the network receives from a given source population *P* by calculating

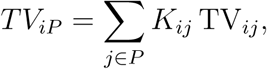

where *K*_*ij*_ = 1 if neuron *j* makes a synapse onto neuron *i*, and *K*_*ij*_ = 0 otherwise.

### Detailed model descriptions

In the following, the mathematical model implementations are described in detail. While this documentation allows to fully reproduce the model, it is not essential for the comprehension of the results of the study. In the main text of the manuscript we do not refer to these details, and the reader may choose to directly jump to **Results**.

#### Model A: Spiking neural network model

We start by describing the spiking neural network model in detail. To a large extent, it is identical to the original model introduced by Potjans and Diesmann (2014). It consists of *N* = 77 169 neurons distributed over eight populations, one excitatory and one inhibitory population in each out of four layers, respectively (Fig 1A). The individual population sizes are summarized in Fig 1C and Table 2. We use the linear leaky integrate-and-fire neuron model throughout this study. In this point neuron approximation, the membrane potential *V*_*i*_(*t*) of each individual neuron follows the differential equation

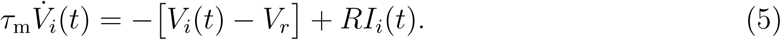

Here, *τ*_m_ is the membrane time constant and *V*_r_ is the resting potential. The total input current has three separate components 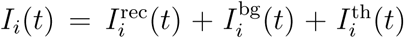. The recurrent input is given by

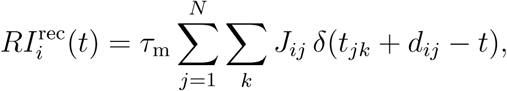

where *t*_*jk*_ is the time of the *k*-th spike of neuron j, *d*_*ij*_ is the transmission delay of the connection, and *J*_*ij*_ is the amplitude of the post-synaptic potential (synaptic “efficacy”) from neuron *j* to neuron *i*. Each time the pre-synaptic neuron *j* fires a spike at time *t*_*jk*_, the membrane potential of neuron *i* is deflected by *J*_*ij*_ at time *t*_*ik*_ + *d*_*ij*_. If the membrane potential reaches the threshold voltage *V*_thr_, the neuron emits a spike that is transmitted to all its post-synaptic neurons. Following each spike, the membrane potential is clamped at the reset potential *V*_*i*_ = *V*_*r*_ for a refractory period *τ*_ref_. In our work, the synapse model slightly differs from the original model of Potjans and Diesmann (2014). While they used exponential post-synaptic currents, in our model all of the charge is delivered instantaneously, as expressed by the *δ*-function in Eq. 5. Numerical values for the single neuron parameters are summarized in Table 1.

**Table 1:**
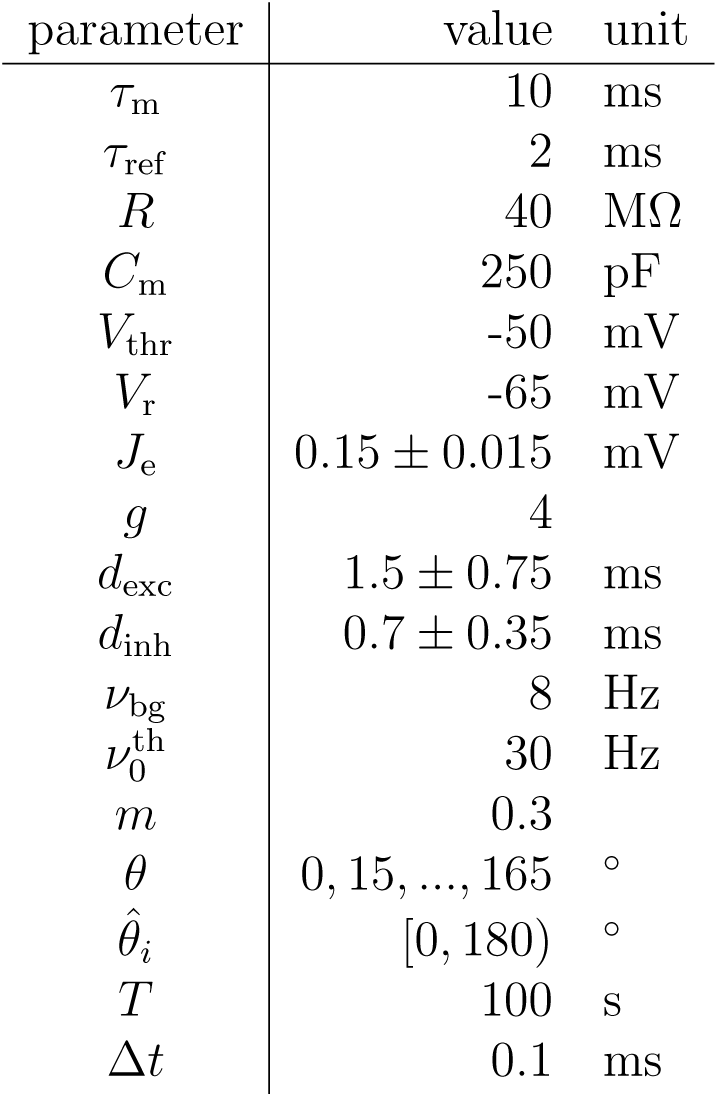
Single neuron parameter values. Shown are variable names, numerical parameter values and, if applicable, the standard deviation of the Gaussian distribution, as well as physical units.

| parameter | value | unit |
| --- | --- | --- |
| $\tau_m$ | 10 | ms |
| $\tau_{\text{ref}}$ | 2 | ms |
| $R$ | 40 | MΩ |
| $C_m$ | 250 | pF |
| $V_{\text{thr}}$ | -50 | mV |
| $V_r$ | -65 | mV |
| $J_e$ | $0.15 \pm 0.015$ | mV |
| $g$ | 4 | |
| $d_{\text{exc}}$ | $1.5 \pm 0.75$ | ms |
| $d_{\text{inh}}$ | $0.7 \pm 0.35$ | ms |
| $\nu_{\text{bg}}$ | 8 | Hz |
| $\nu_0^{\text{th}}$ | 30 | Hz |
| $m$ | 0.3 | |
| $\theta$ | 0, 15, ..., 165 | ° |
| $\hat{\theta}_i$ | [0, 180) | ° |
| $T$ | 100 | s |
| $\Delta t$ | 0.1 | ms |

**Table 2:** Populations sizes.

| population | L2/3e | L2/3i | L4e | L4i | L5e | L5i | L6e | L6i |
| --- | --- | --- | --- | --- | --- | --- | --- | --- |
| size | 20 683 | 5 834 | 21 915 | 5 479 | 4 850 | 1 065 | 14 395 | 2 948 |

The connectivity of the model was compiled in Potjans and Diesmann (2014) based on a large number of anatomical and electrophysiological measurements, most prominently by Thomson et al. (2002) and Binzegger et al. (2004). The connectivity map shown in Fig 1B and Table 3 summarizes the result of an extended review of the literature. In our implementation, these numbers describe the exact number of synapses each post-synaptic neuron receives from randomly chosen pre-synaptic neurons in each of the eight populations. All connections are unitary, and self-connections are excluded. This fixed in-degree connectivity deviates from the original model, where a fixed number of synapses was distributed by randomly drawing both target and source neurons, resulting in approximately binomially distributed in- and out-degrees. Our modification results in a somewhat lower variability of rates and orientation selectivities in the populations.

**Table 3:** In-degrees for individual projections.

|  |  | presynaptic |  |  |  |  |  |  |  |  |  |
| --- | --- | --- | --- | --- | --- | --- | --- | --- | --- | --- | --- |
|  |  | L2/3e | L2/3i | L4e | L4i | L5e | L5i | L6e | L6i | bg | th |
| postsynaptic | L2/3e | 2 199 | 1 079 | 979 | 467 | 159 | 0 | 109 | 0 | 1 600 | 0 |
|  | L2/3i | 2 990 | 860 | 703 | 289 | 380 | 0 | 60 | 0 | 1 500 | 0 |
|  | L4e | 159 | 34 | 1 117 | 794 | 32 | 0 | 667 | 0 | 2100 | 93 |
|  | L4i | 1 480 | 16 | 1 813 | 953 | 16 | 0 | 1 608 | 0 | 1 900 | 57 |
|  | L5e | 2 188 | 374 | 1 135 | 31 | 420 | 496 | 296 | 0 | 2 000 | 0 |
|  | L5i | 1 165 | 159 | 570 | 12 | 300 | 404 | 124 | 0 | 1 900 | 0 |
|  | L6e | 325 | 38 | 467 | 91 | 285 | 21 | 581 | 752 | 2 900 | 47 |
|  | L6i | 766 | 5 | 74 | 2 | 136 | 8 | 979 | 459 | 2 100 | 17 |

Synaptic efficacies *J*_*ij*_ are drawn from a normal distribution with a mean that depends on the type of connection, and a standard deviation corresponding to 10% of the mean. The mean is *J*_e_ for excitatory and *J*_*i*_ = *-g J*_e_ for inhibitory connections. As in the model by Potjans and Diesmann (2014), the projection from L4e to L2/3e neurons is an exception, where a mean efficacy of 2 *J*_e_ is imposed. This modification is due to inconclusive data for this particular projection, see (Feldmeyer et al., 2006; Potjans and Diesmann, 2014) for details. For all unconnected pairs, we set *J*_*ij*_ = 0. In order to obey Dale’s principle, the normal distributions are clipped at zero. Similarly, delays *d*_*ij*_ are also drawn from two normal distributions with fixed means and variances, *d*_exc_ for excitatory and *d*_inh_ for inhibitory connections, respectively. The delay distributions are clipped at the simulation time step Δ*t*. Numerical values for the distributions of connection strengths and delays can be found in Table 1.

In addition to recurrent connections, neurons receive two types of external input. The constant background input represents connections from other brain regions as well as from neurons within V1, which are not part of the model network. It is realized as a homogeneous Poisson input for each neuron. The resulting input current to each single neuron is

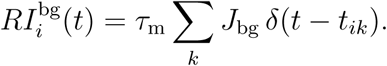

In this case *J*_bg_ = *J*_e_ is the connection strength of background synapses, and *t*_*ik*_ is the time of *k*-th spike of the Poisson process of neuron *i*. The rate of each Poisson process depends on the population *P* the neuron belongs to and is given by 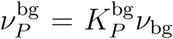. Here, 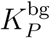 is the number of background synapses connecting to individual neurons in the respective target population (Fig 1B/D and Table 3) and *ν*_bg_ is the rate of individual background neurons.

Similar to background input, the thalamocortical input is also conceived as a homogeneous Poisson process. The current from the connections is given by

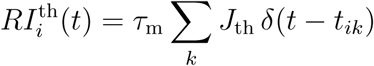

with *J*_th_ = *J*_e_. In contrast to the homogeneous background input to each individual neuron, the rate of the thalamic input depends on the orientation *θ* of a grating that represents the visual stimulus (Fig 1C). The resulting orientation-tuned input is given by

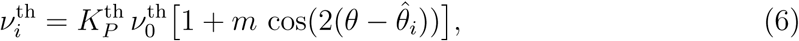

where 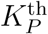 denotes the number of thalamocortical synapses per neuron in population *P* (Fig 1B/E), 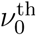 is the fixed mean rate of thalamic neurons, *m* is the modulation strength and 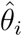 the preferred orientation of the input to this neuron. The latter is drawn from a uniform distribution on [0*°*, 180*°*[. The network is simulated for twelve stimulation angles *θ* from 0*°* to 165*°* in steps of 15*°*. Again, numerical values for all parameters can be found in Table 1 and Table 3.

The non-homogeneous thalamic input is our most significant modification of the model by Potjans and Diesmann (2014) to account for visual stimulation of the network. The original model used a population of 902 thalamic neurons. However, in order to include orientation tuning into the projection, it would be necessary to make assumptions on how the input tuning is generated by the thalamocortical afferents. Because our work concentrates on the cortical processing of tuning information, we replaced the thalamic neurons by Poisson type tuned input.

Stimulation by a grating with each of the twelve angles *θ* is simulated for a total of *T* = 100 s, with a simulation time step of Δ*t* = 0.1 ms. In addition, before the stimulation starts, the network is simulated for a warm-up time of 200 ms to exclude onset transients from the analysis. Spike times and the exact synaptic connectivity for all pairs of neurons are extracted and stored for further analysis.

#### Model B: nonlinear firing rate model

To derive a firing rate model for the system under study, we apply the mean field approach put forward by Brunel (2000) to each single neuron in all eight populations. We assume that the total input to each neuron *i* amounts to a mean current *µ*_*i*_ and additive current fluctuations (Gaussian white noise) *η*(*t*) with variance 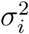

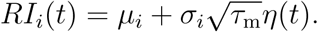

This way, we can characterize the stationary state, in which *µ*_*i*_ and *σ*_*i*_ are fixed parameters that depend on the thalamic input. Similar to the total input current, the mean and variance can be separated into three sources

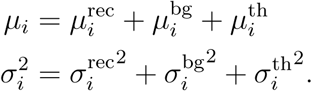

For each source, the respective mean and variance are approximately given by

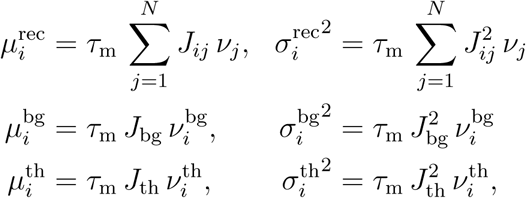

where independence of all input components has been assumed. As for the spiking model, the in-degrees of the external input has been accounted for in 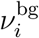 and 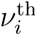. Furthermore, 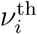 accounts for the non-homogeneous input each individual neuron receives due to the oriented stimulus according to Eq. 6. The self-consistent solutions of the associated first-passage time problem (Siegert, 1951) yield the steady-state firing rates of all neurons

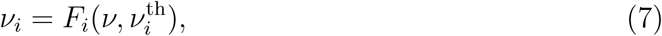

with the transfer function *F*_*i*_ being defined by

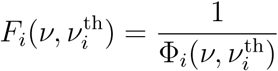

Where

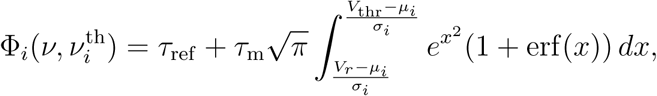

where erf is the standard error function. Solving this non-linear algebraic set of equations yields the desired estimates for the firing rate of each single neuron.

The challenge is now to actually solve this 77 169-dimensional system of highly non-linear self-consistency equations. This can be achieved by re-formulating the problem as a Wilson-Cowan type model (Wilson and Cowan, 1972). Since we are only interested in the fixed point describing the steady-state of the system, and not in its exact temporal evolution, we choose a unit time constant and introduce a pseudo-time *s*. This allows to write the problem as a system of ordinary differential equations

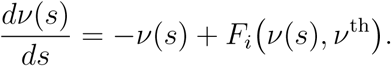

Assuming stability, these differential equations converge to a steady-state rate vector *ν \** which solves the system in Eq. 7 characterized by 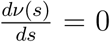. For the numerical solution, we applied an explicit Runge-Kutta method of order (4)5 including adaptive step size control (Hairer et al., 1993).

#### Model C: linearized network model

The linear model is derived from the rate coding model described above using a standard linearization procedure. We start by calculating the total derivative of the transfer function Eq. 7

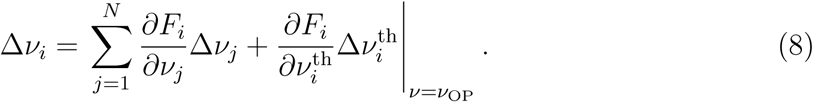

The partial derivatives of the transfer function are given by

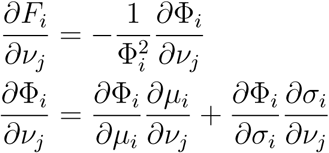

where the partial derivatives of Φ_*i*_ are

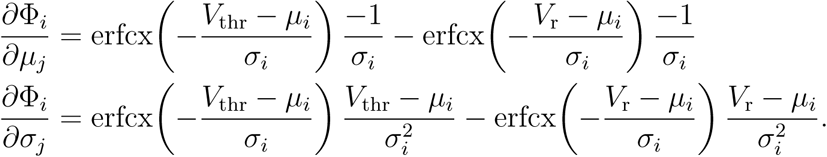

Here, the Leibniz integral rule was used, and 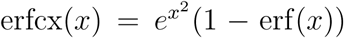 is the scaled complementary error function. Finally, the partial derivatives of the mean and standard deviation of the input are given by

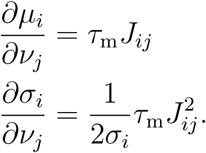

Note that in all partial derivatives above, an explicit mention of the evaluation point *ν* = *ν*_OP_ was omitted for the sake of a compact notation.

The operating point *ν*_OP_ at which the derivatives are evaluated can be chosen freely. However, the precision of the linear approximation depends on the nature of the non-linearity of the activity of the network, and how far the linearization point is away from the regime of interest. Here, it is chosen as the activity with zero thalamic input, *ν*^th^ = 0.

This choice provides the greatest flexibility in the predictions to be derived from the model. In this case, for both conditions considered here, the thalamic input perturbation is identical to the thalamic input rates, 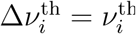. Note that, while thalamic neurons do not fire at the linearization point, the network does still receive background input from other sources, leading to non-zero network activity at this point.

The derivatives of the transfer function are organized in a matrix *W*

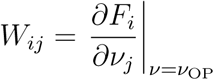

and the diagonal matrix

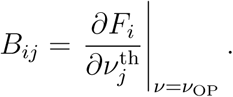

Eq. 8 can then be compactly written as

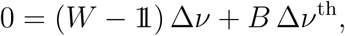

where we also summarized the input and output rate perturbations into the vectors Δ*ν* and Δ*ν*^th^. Assuming that (*W* - 𝟙) is invertible, which is almost always the case for the large random matrices considered here, and defining the scaled effective input Δ*β* = *B* Δ*ν*^th^, we obtain the explicit expression for the output perturbation

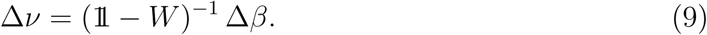

In a next step, the effective input perturbation can be decomposed into

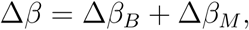

where Δ*β*_*B*_ is the baseline and Δ*β*_*M*_ is the modulation component, respectively (Sadeh et al., 2014). The former is defined by 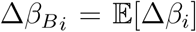, the vector of expected effective input rates. Note that the expectation values are identical across all neurons of each population. With this, we define Δ*β*_*M*_ = Δ*β* - Δ*β*_*B*_ as the vector containing the deviations from the population means.

Along the same lines, the matrix *W* can also be decomposed into two matrices such that

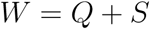

where *Q* is a block-wise constant matrix containing expectation values *Q*_*ij*_ = 𝔼[*W*_*ij*_], and *S* is composed of the zero-mean deviances *S* = *W - Q*. Combining the decomposition of the input perturbation and the matrix, we can define two linear systems

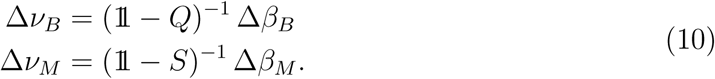

In analogy to what was suggested by Sadeh et al. (2014), these two systems can be interpreted as two separate processing pathways of the network. In the baseline system, the matrix *Q* defines the processing of the untuned baseline input, while in the modulation system, the matrix *S* is responsible for the input modulation. However, this characterizes the system only, if there is no interference between the two pathways: The output Δ*ν*_*B*_ should not contain a modulation component, and the output Δ*ν*_*M*_ should not contain a baseline component. Furthermore, it is not obvious that the combination Δ*ν* = Δ*ν*_*B*_ + Δ*ν*_*M*_ solves the original *W*-system described in Eq. 9.

We start verifying these requirements by showing that Δ*ν*_*B*_ is constant across populations and thus does not contain a modulation component. To this end, a symmetry argument can be applied: Let 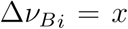 and 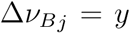 for two neurons *i* and *j* of the same population, then 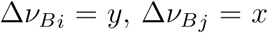 must also be a solution of the system, as *Q* is block-wise constant. As the operator (𝟙*- Q*)^*-*1^ is invertible and the system has only one solution, it follows that *x* = *y* and Δ*ν*_*B*_ must be constant for all neurons from the same population.

In order to show that Δ*ν*_*M*_ does not have a baseline component, we look at the modulation system (Eq. 10) in its element-wise, implicit form

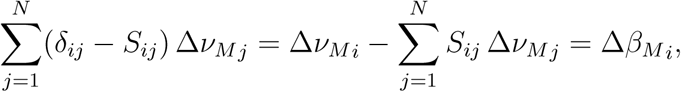

where *d*_*ij*_ is the Kronecker delta function.Changing to the expectation value of the expression and assuming independence of *S*_*ij*_ and 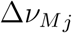 leads to

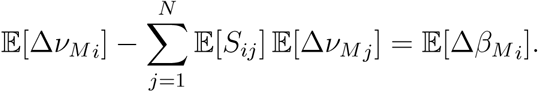

Noting that 𝔼[*S*_*ij*_] = 0 and 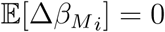 by construction, this shows that

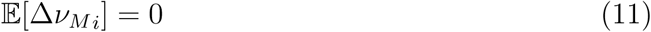

and thus that Δ*ν*_*M*_ has no baseline component.

Finally, we show that Δ*ν* = Δ*ν*_*B*_ + Δ*ν*_*M*_ yields an approximate solution of the *W* - system. For this, we substitute this ansatz and the decomposition *W* = *Q* + *S* into Eq. 9

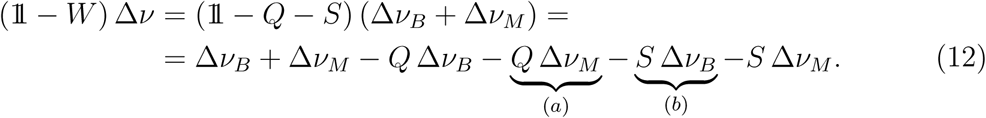

We will now analyze the two interference terms (a) and (b) in more detail

1. The matrix Q is block-wise constant with value *Q*_*ij*_ = *q*_*P T*_ for neurons *i* and *j* in populations *P* and *T*, respectively. For each element in the vector *Q* Δ*ν*_*M*_, we then have

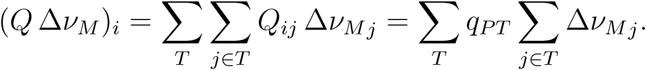

Using Eq. 11, we know that*j∈T* Δ*ν*_*M*_ *≈* 0 and therefore also *Q* Δ*ν*_*M*_ *≈* 0.

2. Similarly, for each element of the second interference term, we can write

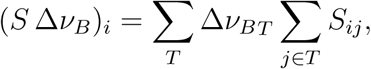

where now 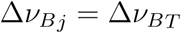 are the values of the population-wise constant vector Δ*ν*_*B*_. The matrix *S* is defined as the zero-mean deviations between *W* and *Q*. Therefore, for sufficiently large population sizes and small variances in the connectivity, we have Σ*j∈T S*_*ij*_ *≈* 0 which leads to *S* Δ*ν*_*B*_ *≈* 0.

Combining these findings with Eq. 12, the expression can be reduced to

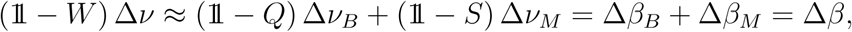

by definition of Δ*ν*_*B*_ and Δ*ν*_*M*_ (Eq. 10), meaning that Δ*ν* = Δ*ν*_*B*_ + Δ*ν*_*M*_ indeed provides an approximate solution to the system.

To summarize, we showed that instead of solving the linear model via the *W* –system directly, it can also be studied in terms of the two pathways defined by Eq. 10. Provided the two interference terms in Eq. 12 can indeed be neglected for the network at hand, this is approximately equivalent and potentially more informative than the direct solution.

### External stimulation

In addition to the synaptic input provided by the background and the thalamus, neurons can also receive additional external stimulation *I*^stim^ in an experiment. Such an input could, for example, model optogenetic microstimulation. In the single neuron rate model (Model B), this can be accounted for by an additional term in the total mean input *µ*_*i*_ given by 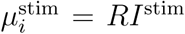. As we consider here only constant stimulation, the variance *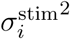* of this input is zero.

In the linear model (Model C), applying a current results in an additional input perturbation and an extra term in the total derivative (Eq. 8). In the explicit expression Eq. 9, this can be accounted for by

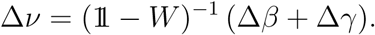

The components of Δγ are defined in analogy to the thalamic perturbation by

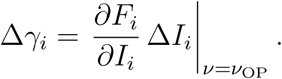

## Results

### Spiking neuronal network simulations

The results of the spiking neural network simulations performed in NEST, and of the detailed data analysis performed on the simulated data, are summarized in Fig 2, both for the spontaneous (weak, untuned thalamic input) and stimulated (tuned thalamic input) condition. Note that we do not subtract the spontaneous rates in the stimulated condition for our analysis.

**Figure 2:**
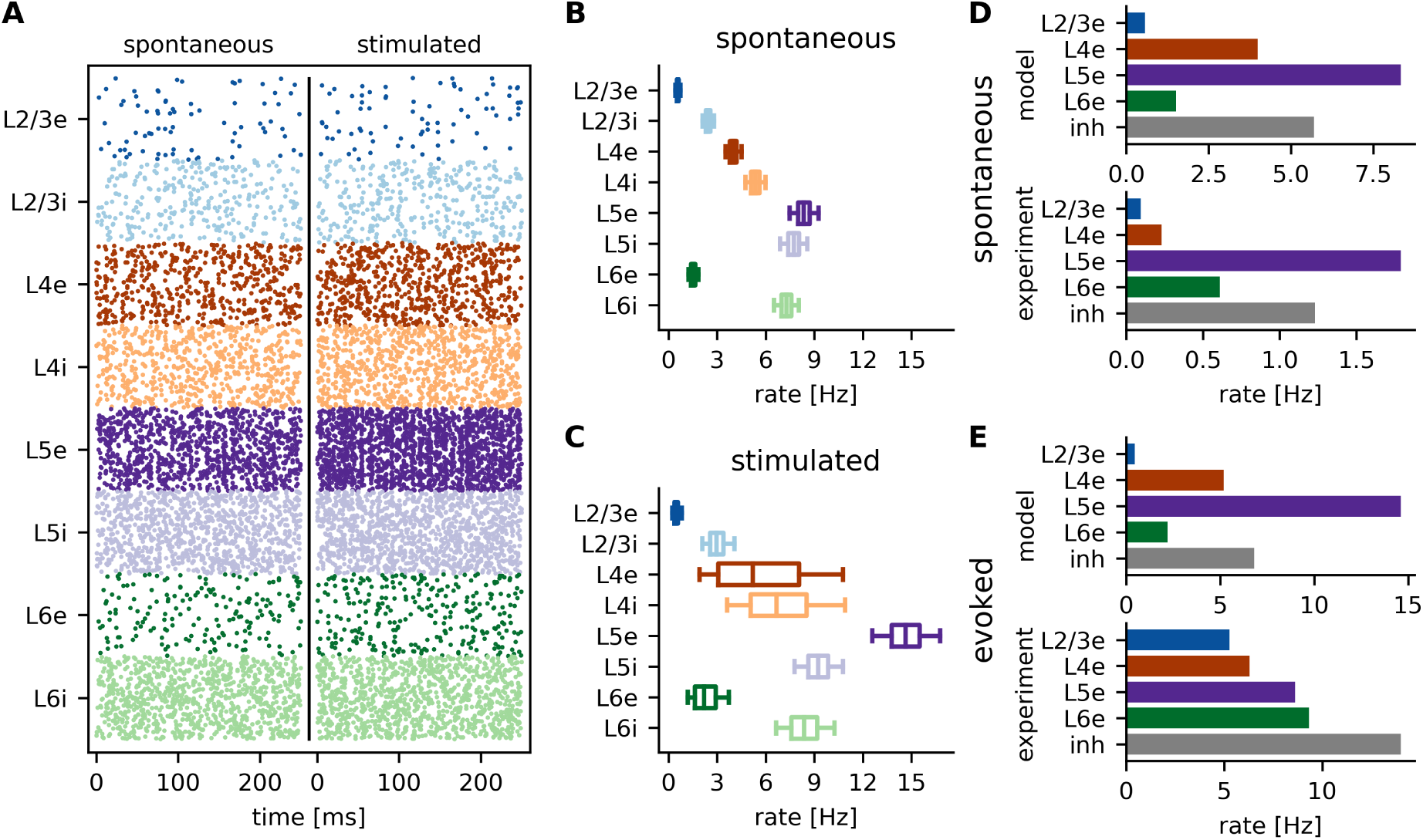
Spiking neuronal network. **A** 250 ms raster plot for both spontaneous and stimulated conditions, depicting 500 neurons of each of the eight subpopulations. **B-C** Box plots of the firing rate distribution within each population, for spontaneous (B) and stimulated (C) conditions, respectively. Shown are the median (central line), lower and upper quartiles (box) as well as 5% and 95% percentiles (whisker bars). **D-E** Comparison of median spontaneous (D) and evoked (E) rates between model results and experimental values reported by (Niell and Stryker, 2008). Evoked rates are given by the maximal response over all angles. The experimental results were obtained by summing the spontaneous and evoked rates from (Niell and Stryker, 2008) for direct comparison. For the model results, the inhibitory rates are averaged over all layers for better comparison with the data from experiments.

As expected, the single neuron firing rates (Fig 2B/C) match the values reported by Potjans and Diesmann (2014). Fig 2D/E compares the firing rates of the model with experimental values obtained by Niell and Stryker (2008) from adult mouse visual cortex. In the spontaneous condition (Fig 2D), the model firing rates are somewhat lower compared to experimental values, but are otherwise in good qualitative agreement with them. We find very low activity in L2/3 and higher rates in the granular and infragranular layers, with the exception of population L6e, where rates are also low.

For the stimulus condition, Fig 2E compares the evoked rates of the model with the results of Niell and Stryker (2008). The evoked rates are defined as the maximal rate of the individual neurons over all angles. Therefore, they differ from the median values shown in Fig 2C. In this condition, the model rates vary more strongly across layers as compared to the experimental values.

The mean and standard deviation (SD) of the pairwise spike-count correlations over all populations are (0.2 *±* 10.3)·10^*-*3^ and (0.3 *±* 10.4)·10^*-*3^ for spontaneous and stimulated conditions, respectively (10 ms bin size). The mean and SD of the coefficient of variation (CV) of the inter-spike intervals are 0.9 *±* 0.1 for both conditions. Both measures show only small variations over different populations (see SFig 1 and SFig 2 for population-wise quantification). The low correlation and Poisson-like irregularity indicate that the network indeed operates in the asynchronous irregular regime (Brunel, 2000; Renart et al., 2010), as illustrated by the raster plot in Fig 2A.

The activity of the network is robust to changes in important parameters which are not well constrained by experiments. Although average firing rates vary across parameter sets, their distribution remained qualitatively similar (SFig 4-9A/B). Only for very low background input, the network operates in a more synchronous regime with increased correlations (SFig 3B/C/SFig 6D). In this case, the network activity also changes qualitatively. Surprisingly, this is only the case for reduced input. For a two-fold increase, the activity remains stable (SFig 3B/SFig 5D)

Our model extends the work of Potjans and Diesmann by giving the thalamic input a weak orientation bias, as described previously (Hansel and van Vreeswijk, 2012; Sadeh et al., 2014). The macroscopic connectivity pattern of the network (Fig 1A) already gives a rough idea of the signal flow through the network. Thalamic afferents project mainly to L4 and to a smaller extent also to L6. From L4, the signal is transmitted to L2/3, which is thought to be the central processing unit of the microcircuit. Signals are then forwarded to higher brain areas as well as to L5, which is the major output unit for projections to sub-cortical brain areas.

In order to dissect the computations performed by the network, we extract tuning curves of all neurons in the model network. As expected from previous studies (Hansel and van Vreeswijk, 2012; Sadeh et al., 2014; Sadeh and Rotter, 2014, 2015), during stimulation, the weak orientation bias of the thalamic input is amplified to strongly orientation selective responses in the input population (Fig 3). Interestingly, L2/3e also shows strong tuning although these neurons only receive input via random connections from L4 neurons. This phenomenon was previously described by Hansel and van Vreeswijk (2012), where projections from L4e to L2/3 were studied independently of other parts of the network. The strong tuning in L2/3e can be explained by a recurrent tuning amplification in inhibition dominated networks (Hansel and van Vreeswijk, 2012; Sadeh et al., 2014). This effect is enough to create strong output tuning from a small bias in the input, which in turn is due to random sampling of preferred orientations in L4. In this context it may seem even more surprising that orientation selectivity of L5 neurons appears to be weaker compared to the other layers. This finding, which is a direct consequence of the underlying experimentally determined connectivity, is consistent with several measurements of orientation selectivity performed in different rodent species (Heimel et al., 2005; Girman et al., 1999; Niell and Stryker, 2008; Sun et al., 2016; Durand et al., 2016) (Fig 4D). However, it cannot be easily explained by visual inspection of the connectivity matrix. Even though synaptic connectivity is stronger for the projection to L2/3e, L5e neurons also receive a large fraction of their input from L4e (Fig 1B, Table 3). Therefore, as the output tuning of L2/3e neurons is essentially inherited from these neurons, one might expect a stronger tuning also in L5e.

**Figure 3:**
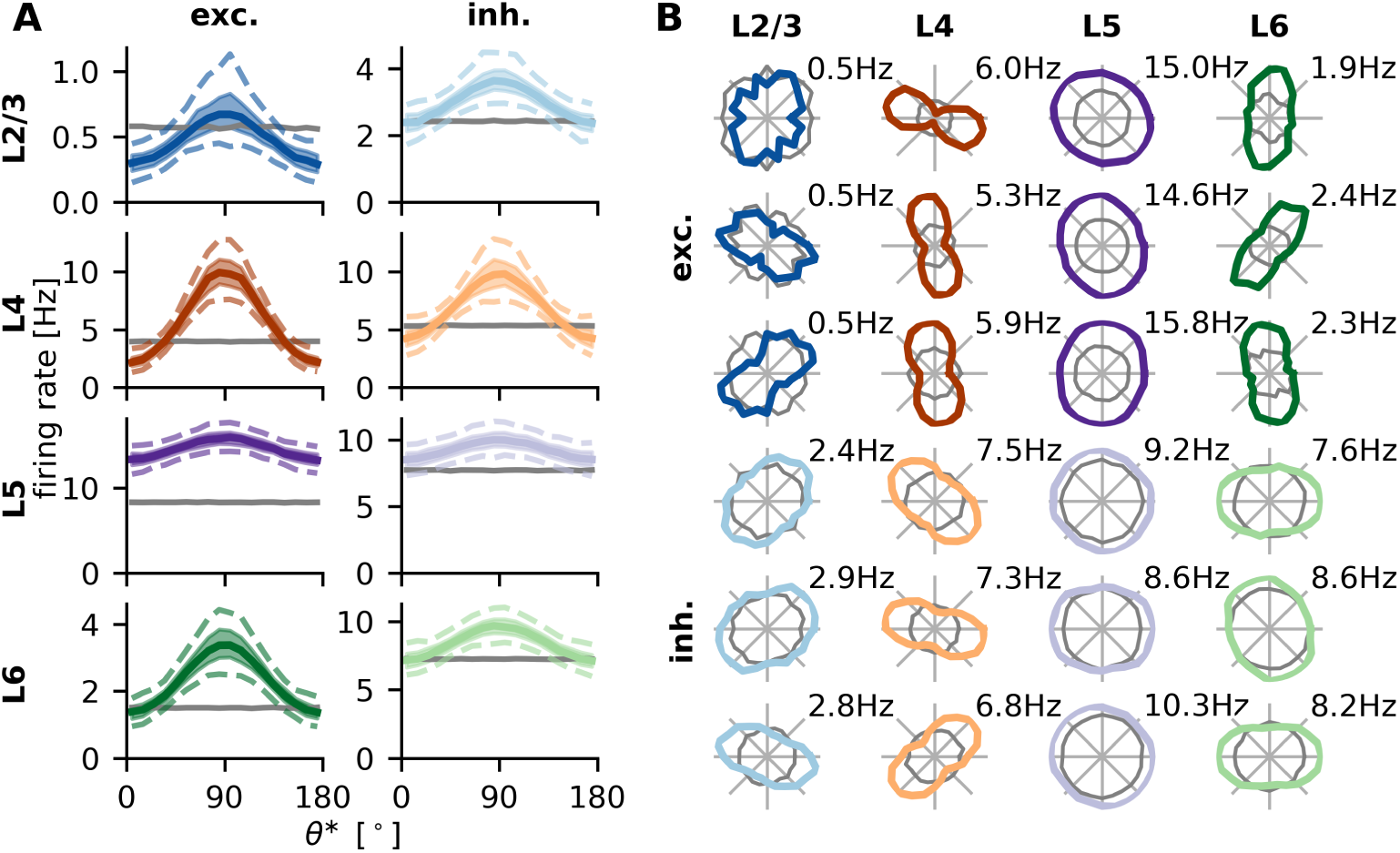
Single neuron tuning curves. **A** Single neuron tuning curves are centered at their respective preferred orientation and then binned over angles (bin size 10*°*). 5%, 25%, 50% (median), 75% and 95% percentiles are then calculated for each bin independently and plotted using different line styles. Gray lines indicate median firing rates in the spontaneous condition. **B** Sample tuning curves from each population in polar representation. Gray lines indicate rates of the same neuron in the spontaneous condition.

**Figure 4:**
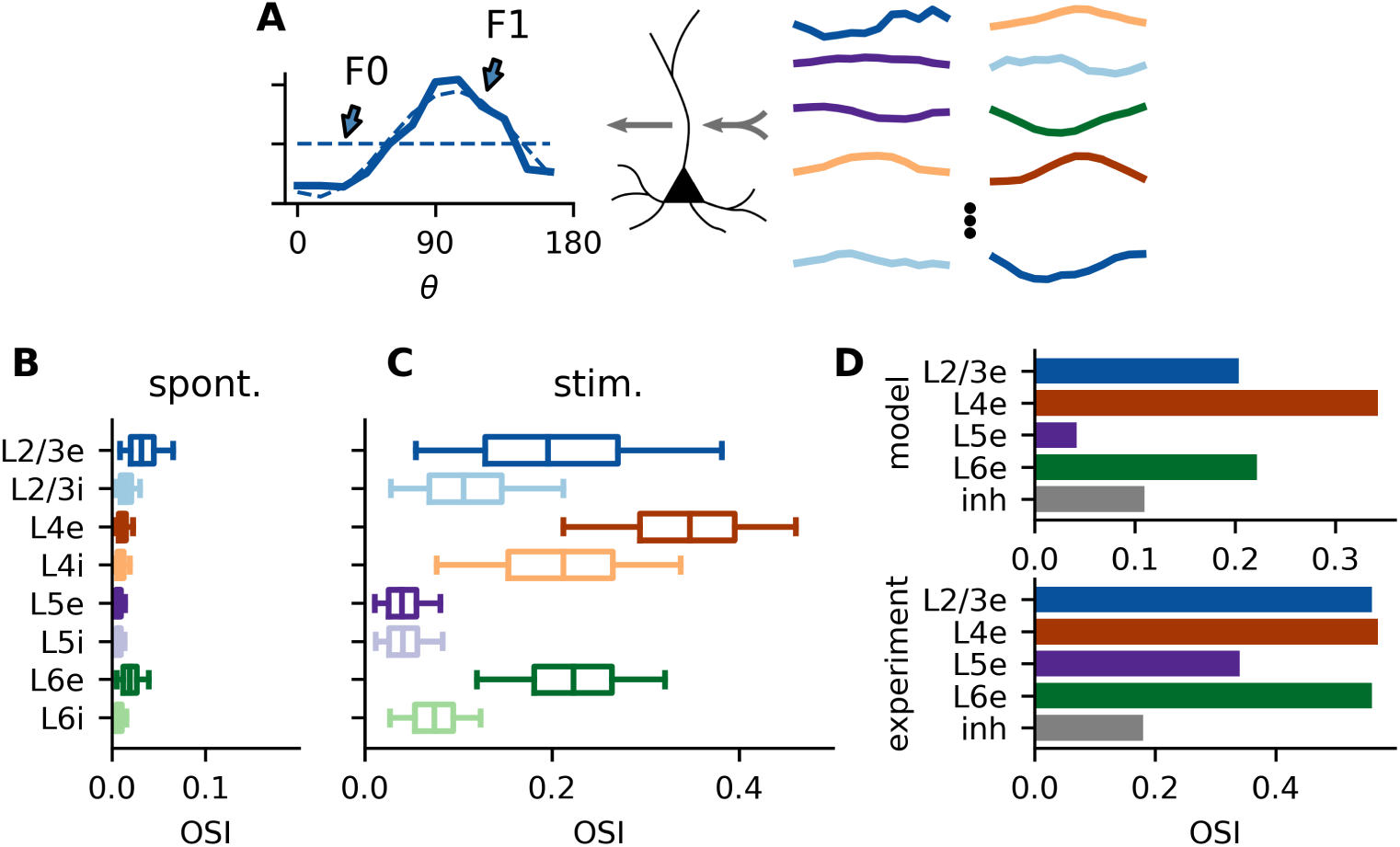
Orientation selectivity in the spiking neural network. **A** Every single neuron integrates its thalamic input with a large number of differently tuned inputs from various cortical populations (right) and generates tuned output from it (left). The orientation selectivity index (OSI) of the neuron is measured by the relative strength of the first (F1) as compared to the zeroth (F0) Fourier component of the output tuning curve (circular variance). **B-C** OSI quantified for neurons in all layers, measured both under spontaneous and stimulated conditions. Percentiles are the same as in Fig 2. **D** Comparison of mean orientation selectivities in the model with experimental values from (Niell and Stryker, 2008).

In order to understand in more detail the differences in orientation selectivity between layers, we extracted the orientation selectivity index (OSI) of all neurons in the model network (Fig 4B/C). We employ a measure based on the circular variance to quantify orientation selectivity (see **Methods** for details). For each neuron, the OSI is calculated as the ratio of first (F1) and zeroth (F0) Fourier component of the tuning curve (Fig 4A). A stronger modulation of the firing rate over angles (higher F1), leads to a higher OSI, while a higher mean rate (higher F0) reduces the OSI. Experimental studies often use a different measure for orientation selectivity based on the activity at the preferred and orthogonal orientation. To allow direct comparison with these works, we also calculated this alternative quantity (SFig 10). While the absolute numbers are generally higher for this measure, the qualitative results are identical. Note that also the distribution of orientation tuning is quite robust against variations in parameters (SFig 4-9C).

Consistent with experimental findings and throughout all layers, inhibitory neurons have a lower OSI than the corresponding excitatory neurons (Niell and Stryker, 2008; Scholl et al., 2015; Durand et al., 2016) (Fig 4D). While this might be expected for L4 and L6 due to the lower number of thalamic afferents, it is not at all obvious for L2/3. In this population, it is a consequence of the recurrent network connectivity, as our analysis will show.

Note that during spontaneous activity, neurons also exhibit weak apparent orientation selectivity (Fig 4B). This is essentially due to fluctuations in the spiking process. Neurons may randomly fire a few action potentials more for one orientation as compared to another, which results in weak but nonzero orientation selectivity. The magnitudes found here are consistent with a Poisson process with the same rate and recording time (data not shown).

The orientation selectivity is measured as the ratio of first and zeroth Fourier component of the tuning curves (Fig 4A). Analyzing the tuning curves in Fig 3A in more detail, we see that although the F1 component in L5e is stronger than in L2/3e, this is overcompensated by the high F0 component of this population, leading to weaker tuning in L5e.

This raises two questions: First, which projections onto L2/3e and L5e lead to the observed difference in activity? Second, why is the tuned (F1) component of L5e not as strongly amplified as the mean rate (F0), similar to the situation in L2/3? In the following, we will address these questions, employing suitable mathematical and computational methods. While the first question can be answered by analyzing the input currents received by the different populations, the second question requires an approach, which takes the strongly recurrent nature of the circuit across layers into consideration.

### Operating point of the network

Each neuron in the eight populations of the network receives inputs from several presynaptic populations, possibly all with different tuning curves, and forms its own output tuning from those (Fig 4A). It has been shown in experiments that this scenario also reflects the situation in rodents (Jia et al., 2010). Which projections are most potent for driving the the target neuron is not immediately obvious from the anatomical connectivity between neurons (Fig 1B). The activity of pre-synaptic neurons is of course also relevant for the total input current to a given neuron. In Fig 5A, the mean input current (Eq. 3) for each projection between populations is shown. While the difference between inputs to L2/3e and L5e seems insignificant when looking at the underlying connectivity (Fig 1B), the picture changes completely if the activity of pre-synaptic neurons is taken into consideration (Fig 5A). The input to L2/3e neurons is mostly determined by L2/3i, both L4 populations and the constant background. Input to L5e, on the other hand, is dominated by L5i and background input, but only to a lesser extent by inputs from L2/3i, L4e and L5e. As a result, due to the different excitation-inhibition (EI) balance, the net currents imply a higher mean input to L5e compared to L2/3e (Fig 5C).

**Figure 5:**
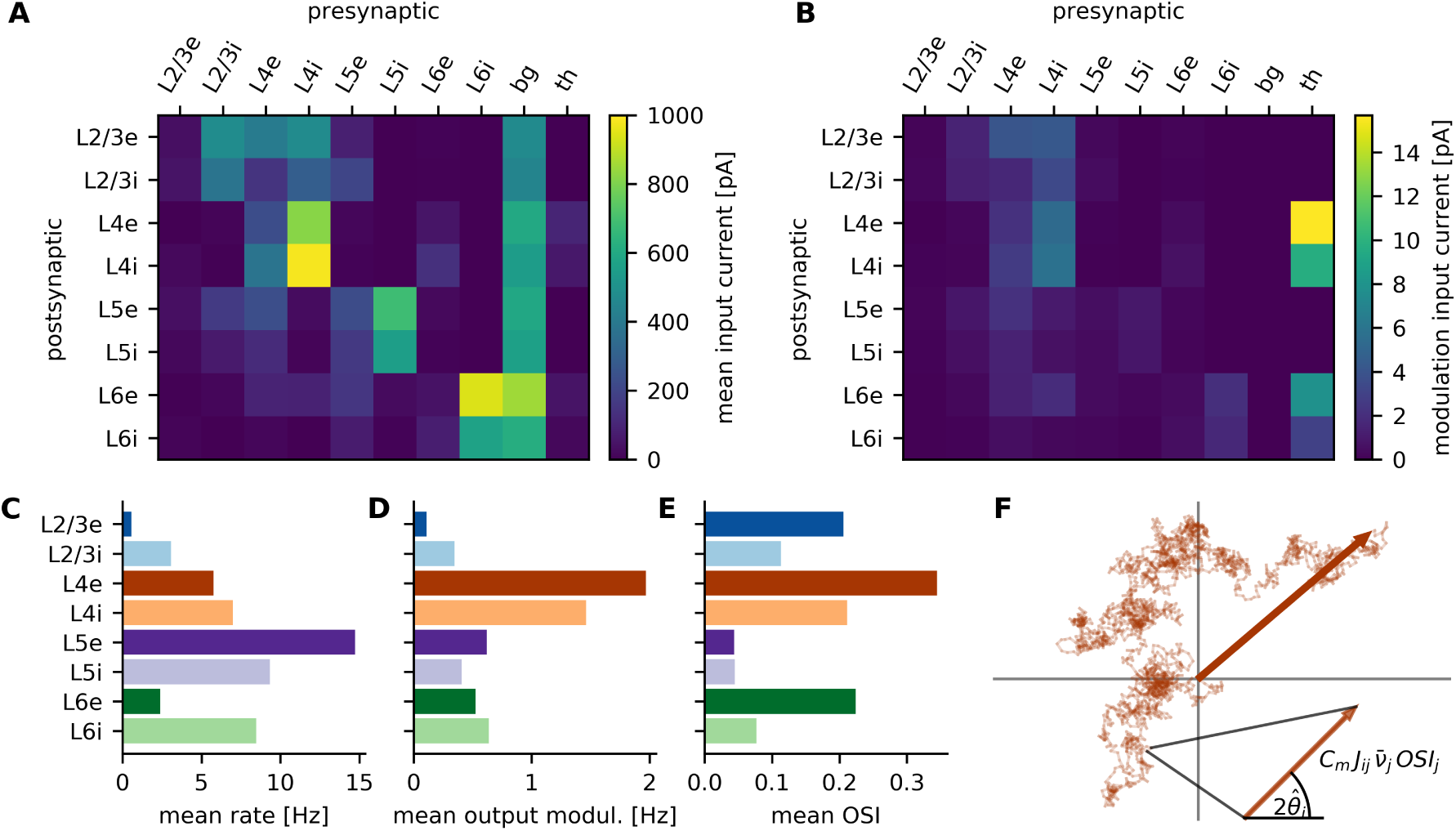
Neuronal input analysis. **A** Mean input currents for all projections between all the involved neuronal populations. **B** Mean projection tuning vector length for all recurrent projections (see text for explanation). **C-E** Output firing rates (F0 of tuning curves), output modulations (F1 of tuning curves) and resulting OSI, averaged over each population. **F** Illustration of single neuron input tuning vector addition. Each small arrow represents the tuning vector of a single pre-synaptic neuron. Linearly adding the vectors of all pre-synaptic neurons of one population results in the tuning vector that characterizes the projection to one post-synaptic neuron.

The marked difference between anatomically defined connectivity (Fig 1B) and the total input current in the stationary balanced state (Fig 5A) highlights the contribution of activity dynamics in recurrent networks (Van Vreeswijk and Sompolinsky, 1996, 1998; Brunel, 2000). While the situation for L5e and L2/3e is very similar when counting the number of excitatory and inhibitory synapses that terminate in each population, the actual current drive they receive is quite different. Since the activity of the whole network autonomously settles in an operating point where the EI balance is dynamically maintained, this can have very different consequences for the various populations, irrespective of the anatomical connectivity.

Having demonstrated how the dynamic equilibrium and the resulting operating point of the layered network explains the high firing rate of L5 neurons, we can now ask the same question with regard to orientation tuning in each neuronal population. In combination, these two aspects fully determine the orientation selectivity index (OSI) of all neurons. To achieve this, we calculate for each neuron a tuning vector summarizing the information about the stimulus conveyed by the input currents to that neuron. Its magnitude measures the tuning strength of the combined input to the post-synaptic neuron, whereas its direction indicates its preferred orientation (Fig 5F inset, Eq. 4). The total tuning input from one specific source population to a single neuron can then be calculated as the sum of all tuning vectors of neurons in the pre-synaptic population which connect to the post-synaptic neuron. If all tuning curves are cosine-like, this concept exactly corresponds to linear summation of inputs. In any case are the magnitudes of the tuning vectors identical to the first Fourier component (F1) of the tuning curves for the current input.

Fig 5B summarizes the mean lengths of all projection tuning vectors, summarizing the information flow in the system. The tuning information enters the cortical network in L4 and L6 and then spreads to the other populations. Although both L4 and L6 neurons show significant tuning to stimulus orientation, due to the low firing rates of L6e neurons it is mostly L4, which provides tuning information to the other populations. In contrast to what would be expected from the connectivity alone (Fig 1B), also L5e neurons receive most of the tuned current from L4. Comparing the input to L2/3e and L5e neurons in more detail reveals that L2/3e neurons mostly integrate tuned input from both populations in L4, while L5e neurons receive less tuned input almost exclusively from L4e.

Considering only the output modulation of the different populations (Fig 5D), which is defined as the first Fourier component of the tuning curves (Fig 4A), it surprises that despite the less tuned input to L5e, these neurons still show a stronger output modulation than L2/3e. This highlights that, besides the input strength, the input sensitivity of neurons is relevant as well. However, the large modulation component of the output of L5e neurons cannot compensate their high firing rates, leading to a lower OSI (Fig 5C-E). Comparing the distribution of firing rates and output modulations over the eight populations (Fig 5C/D), it is evident that these two components are subject to different gain factors. In the following, these dependencies will be studied in more detail.

### Two processing pathways: baseline and modulation

Our analyses so far disentangled the input to neurons in different populations, which explain the observed features of neuronal output. However, the multi-population model is highly recurrent. Therefore, it is not yet clear why the input to all neurons settled in the observed operating point. We face a chicken-and-egg problem here, as in the recurrent network, the output of all populations is simultaneously the input to the same populations, eventually settling in the dynamical operating point. In order to tackle this problem, we now change our perspective from analyzing the input-output relation of single neurons to input-output behavior of the entire network. The idea is to manage the problem by considering the network as a distributed system, which can perform its function only as a whole.

This approach is best realized by linearizing the network dynamics about its operating point. We obtain our close to 80 000-dimensional linearized network model (Model C) by first formulating a single neuron firing rate model based on the single neuron transfer function *F*_*i*_(*ν, ν*^th^) (Model B). The firing rate model is then linearized about its dynamical operating point, leading to the explicit input-output relation

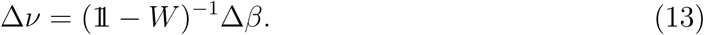

Here, Δ*ν* is a vector summarizing all single neuron rate changes due to a change in the effective input Δ*β*, which in turn is the change in thalamic input rate scaled by the input sensitivities of the different populations. These sensitivities can also be understood as the feed-forward gains of the system (Sadeh et al., 2014), see **Methods** for details. Furthermore, the behavior of the recurrent model is governed by the matrix *W*, which is the matrix of effective recurrent connections, which are a product of the anatomically and physiologically defined synaptic weights and the input-output sensitivities (gains) of individual post-synaptic neurons at their operating point. In combination with the activity at the operating point *ν*_0_, the rate change Δ*ν* results in the output activity *ν* = *ν*_0_ + Δ*ν* of the linear network model.

The explicit form of Eq. 13 also supports the idea of considering the network as an integrated system with one single vector-valued input-output relation. Importantly, this relation is governed by the inverse of the effective connectivity 𝟙 *-W*. Because the entries in *W*_*ij*_ are a function of *K*_*ij*_, this inversion potentially distributes the influence of each single connectivity parameter *K*_*ij*_ over the entire network. This observation also explains why the effects of individual connectivity parameters can be counterintuitive (Tsodyks et al., 1997).

The firing rate model (Model B) as well as the linear network model (Model C) can be considered as simplifications of the spiking neural network (Model A), as they make additional assumptions about its dynamics. Therefore, before analyzing these models in more detail, it is important to establish the consistency of their respective behavior. SFig 11 compares the results from a simulation of all three models, for matched network parameters. We found that the single neuron firing rates of the three models are in good agreement. While the similarity between the non-linear rate model (Model B) and the linear model (Model C) is quite high, both exhibit mild discrepancies from the spiking network model (Model A). In particular, for the larger firing rates of L5e, the non-linear rate model and the linear model tend to slightly underestimate them. Generally, also the preferred orientations and orientation selectivities are in good agreement between the different models. While the match is again excellent for the non-linear firing rate model and linear network model, both slighly deviate from the spiking network model. This can be explained by the role played by activity fluctuations in this model. In particular, the observed orientation selectivities are somewhat higher here, reflecting the same positive bias as the one observed during spontaneous activity.

The match between the three models is also very robust with respect to changes in parameters (SFig 4-9A-C). In fact, even for reduced background input, when there is substantial synchrony in the spiking activity, the non-linear rate and linear models still provide reasonable approximations (SFig 6A-C).

Now that the general consistency in the behavior of the different models is established, we can analyze the linearized network and be confident that our conclusions are also valid for the spiking network. In previous work, a separation into two separate, non-interfering pathways were postulated, exploiting that the network can have different gains for different components of its input (Sadeh et al., 2014) (Fig 6A). Specifically, the effective input perturbation is split into a baseline and a modulation component such that Δ*β* = Δ*β*_*B*_ + Δ*β*_*M*_. Here, we generalize this decomposition to our present multi-population model. Furthermore, we present a novel analytical approach for their analysis.

**Figure 6:**
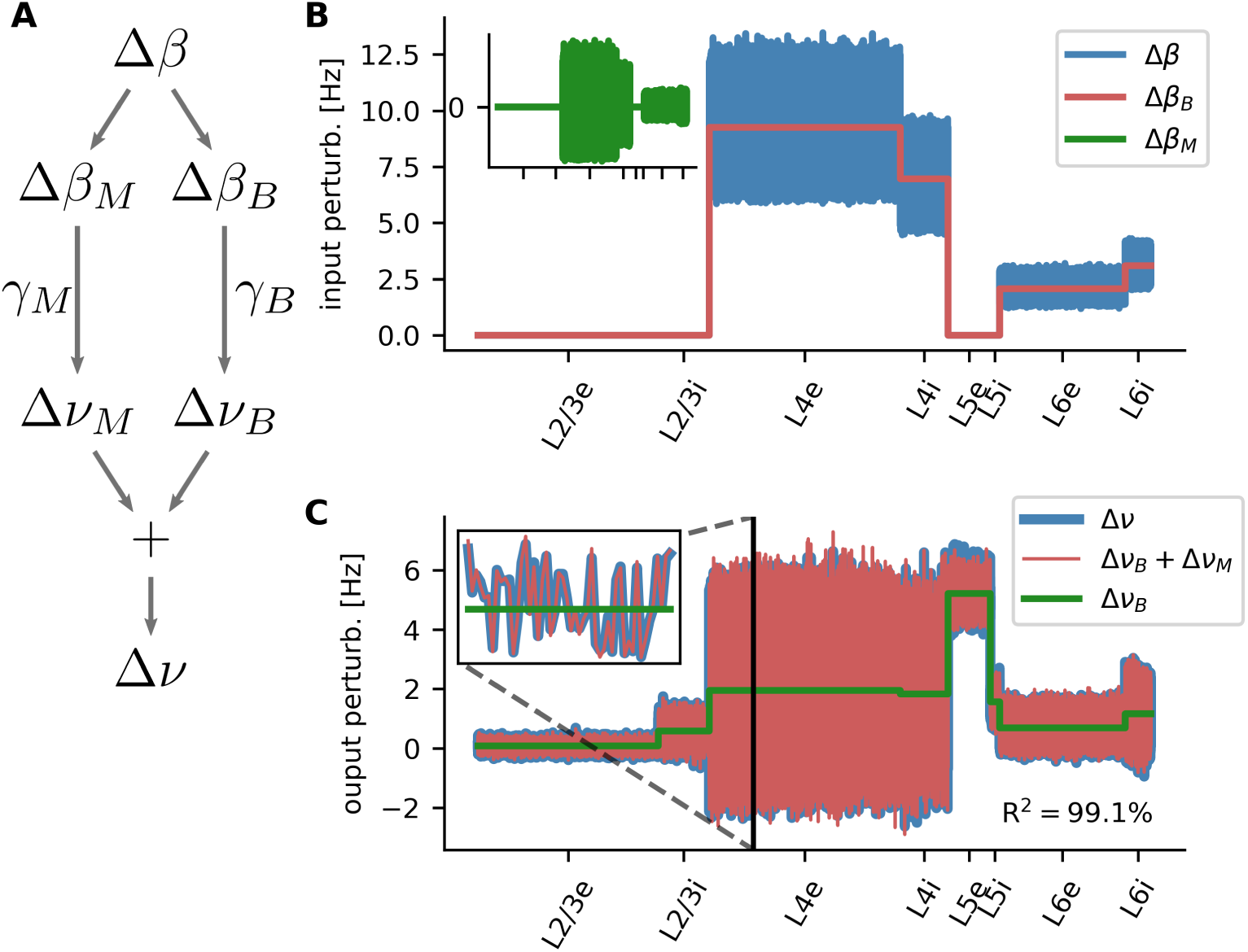
Separate amplification/attenuation of non-informative baseline and tuned modulation. **A** The input perturbation Δ*β* can be decomposed into baseline and modulation, which are then amplified or attenuated by the network with different gains. **B** Input perturbation for one particular orientation of the oriented stimulus. The effective change in input rate is shown for each neuron in the eight populations of the layered network (blue). The baseline (population mean) is depicted in red. The modulation, which is the entry-wise difference between the two, is plotted in the inset (green). **C** Solutions for the baseline and modulation system. Shown are the baseline (green) and baseline plus modulation (red) output rates. For comparison, also the solution of the full *W* system is shown (blue). The inset shows a magnification for a small sample of neurons in population L4e.

For each population, the baseline Δ*β*_*B*_ is conceived as the mean input these neurons receive during stimulation. Since the mean input is different for each population, this leads to a population-wise constant input vector (Fig 6B, red curve). The modulation Δ*β*_*M*_, in contrast, accounts for all neuron-by-neuron deviations from the population base-line, Δ*β*_*M*_:= Δ*β* - Δ*β*_*B*_, yielding a vector with zero mean in each of the eight populations (Fig 6B, inset). Most importantly, the modulation component includes the tuning information that each neurons receives, based on the neuron-specific orientation bias of the stimulus. In other words, the deviation of Δ*β*_*M*_ from the baseline is determined by the orientation of the stimulus, as described by Eq. 6.

Note that the baseline and modulation components are both only indirectly related to mean rate (F0) and tuning strength (F1) discussed in the previous section. While the latter were calculated for the tuning curve of each single neuron independently, the baseline and modulation of the input were calculated over all neurons in one population, for a single stimulation angle.

Separation of baseline and modulation pathway means that the two input components Δ*β*_*B*_ and Δ*β*_*M*_ are processed by two independent mechanisms with possibly distinct gains γ_*B*_ and γ_*M*_. As shown in **Methods**, the two pathways are governed by

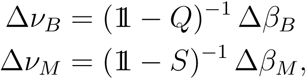

where the decomposition *W* = *Q* + *S* was used. Thereby, *Q* contains population-wise expectation values, resulting in an 8 × 8 block structure, and *S* are the individual modu-lations *S* = *W-Q* of single connections in the network. Importantly, there is no cross-talk between the two pathways. A change in the input baseline Δ*β*_*B*_ does not lead to change in the output modulation Δ*ν*_*M*_, and vice versa.

Solving these two systems independently yields two output rate changes Δ*ν*_*B*_ and Δ*ν*_*M*_. If the cross-terms in the calculations are indeed negligible, the solution Δ*ν* = Δ*ν*_*B*_ + Δ*ν*_*M*_ provides a good approximation to the network behavior (see Eq. 12ff). For the network under study here, the mean magnitudes *µ* of the cross-terms are *µ*[*|Q*Δ*ν*_*M*_ *|*] = 0.02 and *µ*[*|S*Δ*ν*_*B*_ *|*] = 0.09 as compared to *µ*[*|* Δ*β |*] = 3.63, implying that the system can indeed be studied by treating the baseline and the modulation pathway separately. Similar numbers are also obtained for the other parameter sets SFig 3E.

The solutions of the baseline and modulation systems, Δ*ν*_*B*_ and Δ*ν*_*M*_, are shown in Fig 6C in comparison to the solution of the *W*-system. Throughout all populations, the separation solution is in excellent agreement with the direct solution. The modulation part, which conveys the tuning information of the neurons, matches almost perfectly (Fig 6C inset). The good agreement is further confirmed by the high coefficient of determination (“variance explained”) of *R*^2^ = 99.1%. For the different parameter sets, the lowest *R*^2^ is obtained for increased background input with *R*^2^ = 98.0% (SFig 3D/SFig 4-9E). This demonstrates the robustness of the baseline and modulation decomposition also with respect to changes in parameters.

In previous sections, we identified the high baseline rate of L5e as the main cause for the low orientation selectivity of its neurons. This becomes manifest in a high baseline output rate Δ*ν*_*B*_ for this particular population (Fig 6C). While this observation still does not fully reveal the underlying reason for the high rates, it hints at the effective mean connectivity represented by *Q*. In the following, we will therefore study the baseline pathway in more detail.

### Mode decomposition

We start by summarizing the results of Sadeh et al. (2014, 2015), where a similar scenario for a two population EI network was studied. In analogy to the present model, the two population system can also be described by a linear system of the form

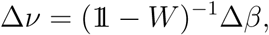

where the dimensionality of the system equals the total number of neurons. In order to study the network behavior, it is instrumental to inspect the eigenvalue spectrum of the matrix *W*. In the two population case, when both populations receive the same input, the eigenvalue spectrum consists of a bulk of known radius localized at the origin and a single exceptional eigenvalue *λ* (Rajan and Abbott, 2006). *For* inhibition dominated networks, this exceptional eigenvalue is real and negative (Fig 7A). Furthermore, the eigenvector Ψ corresponding to that exceptional eigenvalue is the uniform vector, with all entries identical. In this two-population network model, the baseline component of the input perturbation is also proportional to the uniform vector

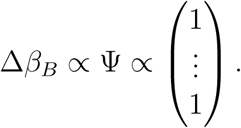

**Figure 7:**
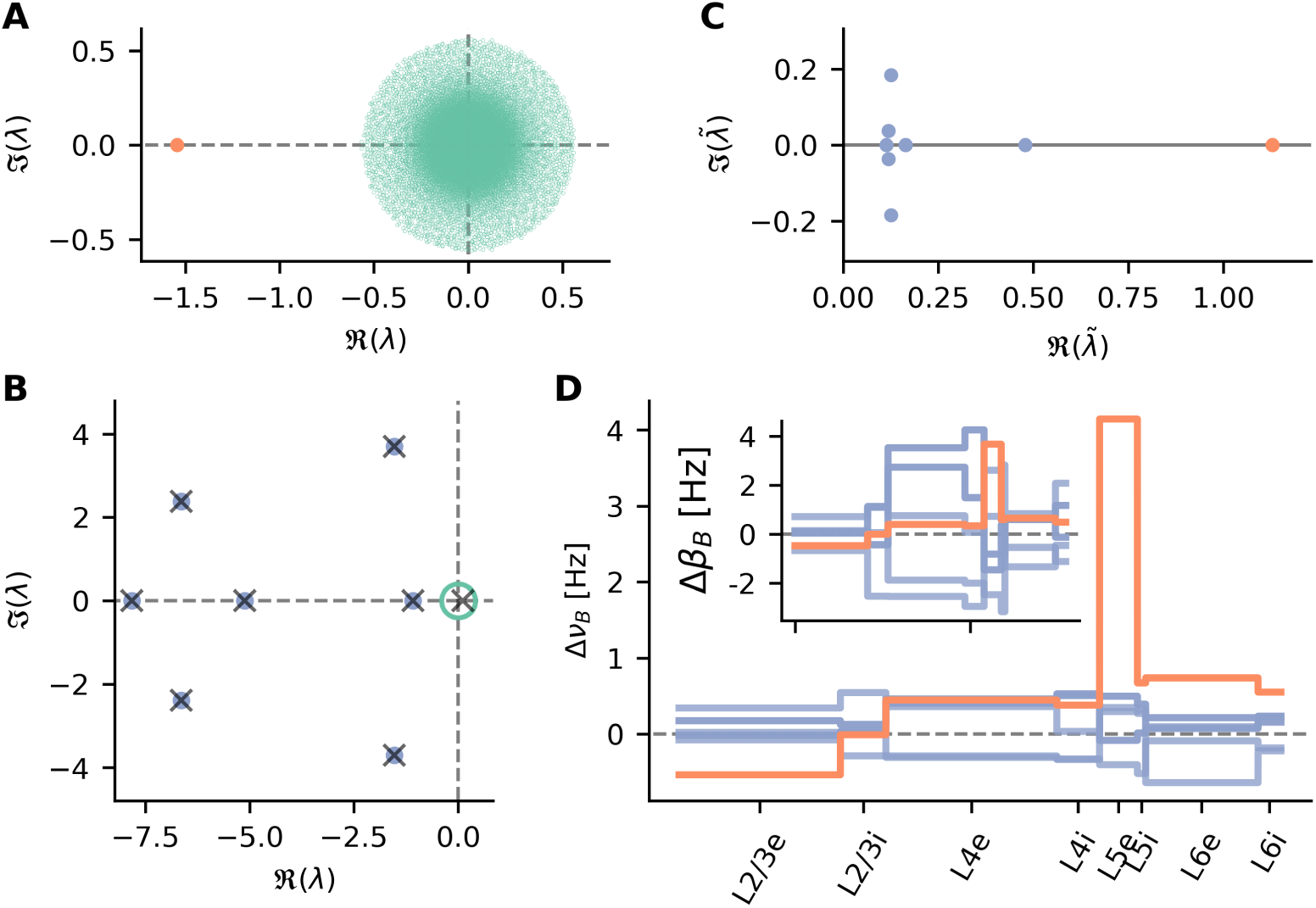
Eigenmode analysis of input-output relations. **A** Eigenvalue spectrum of the linearized model of Brunel (2000). The exceptional eigenvalue is shown in red, the bulk spectrum in green. **B** Eigenvalues of the close to 80 000-dimensional linear system. Blue and green dots indicate the exceptional and bulk eigenvalues of *W*, respectively. Crosses mark eigenvalues of *Q*. **C** Eigenvalues of the matrix (𝟙*- Q*)^*-*1^. **D** The eight output modes of the *Q* system. Shown are the change in output rate for each single neuron, due to each mode. Inset: Input perturbation decomposed into the eight modes, which are also the right eigenvectors of *Q*.

Therefore, exploiting the eigenvector property of a baseline perturbation, the exceptional eigenvalue can be transformed into the gain factor of the baseline

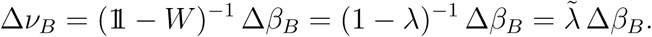

For inhibition dominated networks, this gain factor 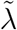 has a small magnitude and thus results in a strong attenuation of the baseline component. This, in turn, amplifies the orientation selectivity of the network.

Generalizing the two-population scenario to the eight-population network considered here, two major differences in terms of the eigenvalue spectrum of *W* become apparent. First, instead of a single exceptional eigenvalue, the spectrum of the effective connectivity is more complex (Fig 7B). In addition to the bulk of eigenvalues (diameter indicated in green in Fig 7B), there are seven eigenvalues with a significantly larger magnitude (blue dots in Fig 7B). In a random matrix theory context, it was previously shown that the exceptional and bulk eigenvalues are due to the expectation values and variances of the distribution of the elements of *W*. Therefore, the exceptional eigenvalues are asymptotically identical to the eigenvalues of *Q* (Aljadeff et al., 2016). *Indeed, the first seven eigenvalues are in good agreement with the numerical calculations on W* (crosses in Fig 7B). Furthermore, an eighth exceptional eigenvalue is identified, which lies in the middle of the bulk and could therefore not be distinguished from bulk eigenvalues in numerical calculations on *W*.

The second difference compared to the two-population scenario is the fact that the constant vector is not any more an eigenvector of an exceptional eigenvalue. Instead, the eigenvectors of the baseline system *Q* are only population-wise constant (cf. Fig 7D inset). However, also the input perturbation is not proportional to the constant vector in this case. For each population, it depends on the sensitivity to thalamic input, also resulting in a population-wise constant function. In order to study the input-output behavior, the input can therefore be decomposed into the eight eigenvectors (cf. Fig 8 left)

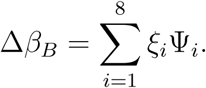

**Figure 8:**
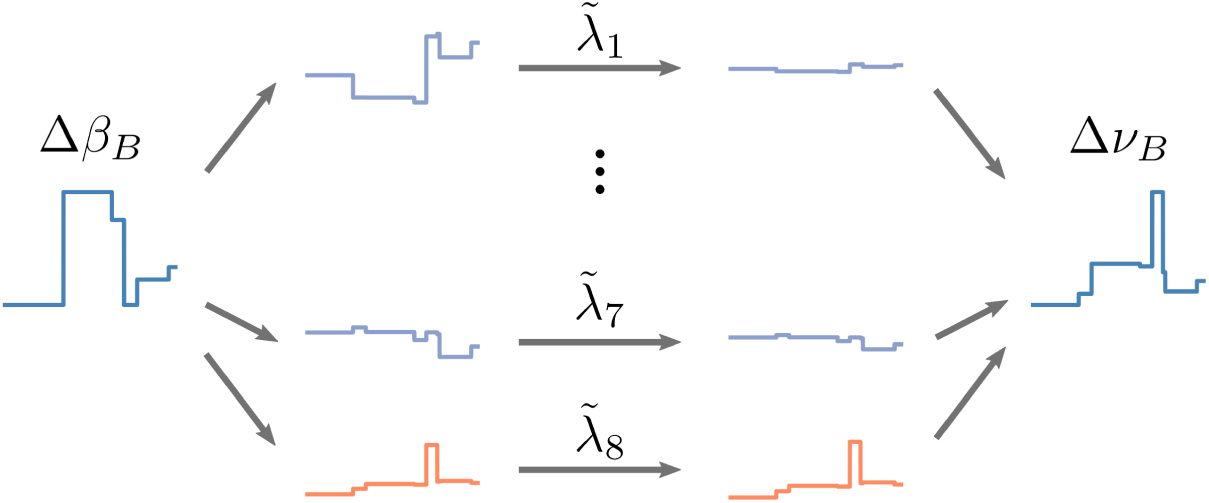
Eigenmode decomposition of input and output. Left: The baseline component of the input perturbation Δ*β*_*B*_ is decomposed into the eight input modes *ξ*_*i*_Ψ_*i*_. Center: By the network action, each mode is amplified or attenuated by its individual gain factor 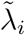. Right: The sum of all scaled output modes 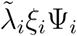 results in the baseline component of the output perturbation Δ*ν*_*B*_.

Here, Ψ_*i*_ are the eigenvectors of *Q* and *ξ*_*i*_ are the coefficients of the modes. The weighted components *ξ*_*i*_Ψ_*i*_ can be considered as the *input modes* of the input perturbation Δ*β*_*B*_. The eight modes which represent the perturbation of the stimulation are shown in the inset of Fig 7D. Note that the sum over these modes is identical to Δ*β*_*B*_ in Fig 6B. Applying the decomposition to the linear baseline system of Eq. 14 results in a decomposition of the output rates of the network, where the eight exceptional eigenvalues define the gain factors of the individual modes by (Fig 8 right)

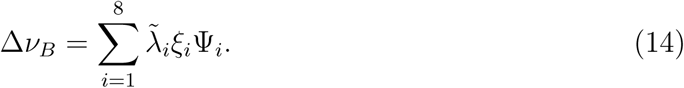

The transformed gain factors 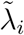 are shown in Fig 7C. The individual parts of this decomposition can be considered as *output modes* of the system, with a direct one-to-one relation to the input modes. The effect of the exceptional eigenvalues *λ* is quite similar to the two-population scenario described previously, where the single exceptional eigenvalue defined the gain of the homogeneous common mode. In the multi-population case, each input mode is amplified with the respective gain 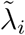 to form the output perturbation of the network. The eight output modes are shown in Fig 7D. Note that, similar to the input perturbation Δ*β*_*B*_, the sum of the modes is identical to the baseline component Δ*ν*_*B*_ shown in Fig 6C.

Studying the output modes in more detail, it is apparent that the high rates of L5e neurons, and thus also their low orientation selectivity, are mostly due to one specific mode (orange line in Fig 7D). On the other hand, the same mode has a similar strength as the other components in the input space (orange line, Fig 7D inset). As the corresponding eigenvalue of *W* has a very small magnitude, however, the gain factor 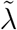 of that mode is much larger compared to the other modes (Fig 7C, orange dot). Due to this gain, the mode is dominant in the output rate change, resulting in high firing rates in L5e. The fact that only a single mode is responsible for this strong amplification underlines our conclusion that this effect is a feature of the whole network, which cannot be pinned down to one specific connection in a meaningful way. Importantly, although the eigenvalue spectrum and mode structure can change to some degree, the observation that a single mode with a strong gain is responsible for the high rates of L5e is consistent for all parameter sets considered (SFig 4-9F/G).

### Predictions for new stimulation experiments

Having identified the large gain of a specific mode in the input as the source of the high firing rates in L5e, we now apply our theory to predict the network behavior for a scenario, where that particular mode is absent in the input. For two reasons the mode cannot be directly removed from the input firing rates. First, that mode has nonzero coefficients for all populations, which leads to non-zero thalamic input also to L2/3 and L5. While it is technically possible to implement this in a computational model, it is infeasible as a biological experiment. The second problem is more fundamental: Subtracting the mode from the input perturbation would lead to negative firing rates, which cannot be realized. Therefore, instead of using altered Poisson input to neurons, we designed an external current stimulation, which has an equivalent effect on the neuronal output rates. It is conceivable to realize such an input, for example by specific optogenetic stimulation. In the linear model (Model C), an external current results in an additional input perturbation Δγ such that

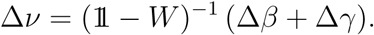

In order to delete a specific mode Ψ_*k*_ from the input, we can set Δ*γ* = *-ξ*_*k*_Ψ_*k*_. As a consequence, when the two inputs Δ*β* and Δγ are combined, the mode cancels out, resulting in altered output perturbation.

The required input current for single neurons in each population are shown in Fig 9A. They are not directly proportional to the deleted mode, due to the scaling by the individual feed-forward gain of each population, as well as due to the highly recurrent processing of the network. As expected, since the rate of L5e neurons should be suppressed, these neurons receive a strongly negative current. More surprisingly, L2/3e requires a positive input current of similar magnitude, although the desired change in rate of that population is much smaller.

**Figure 9:**
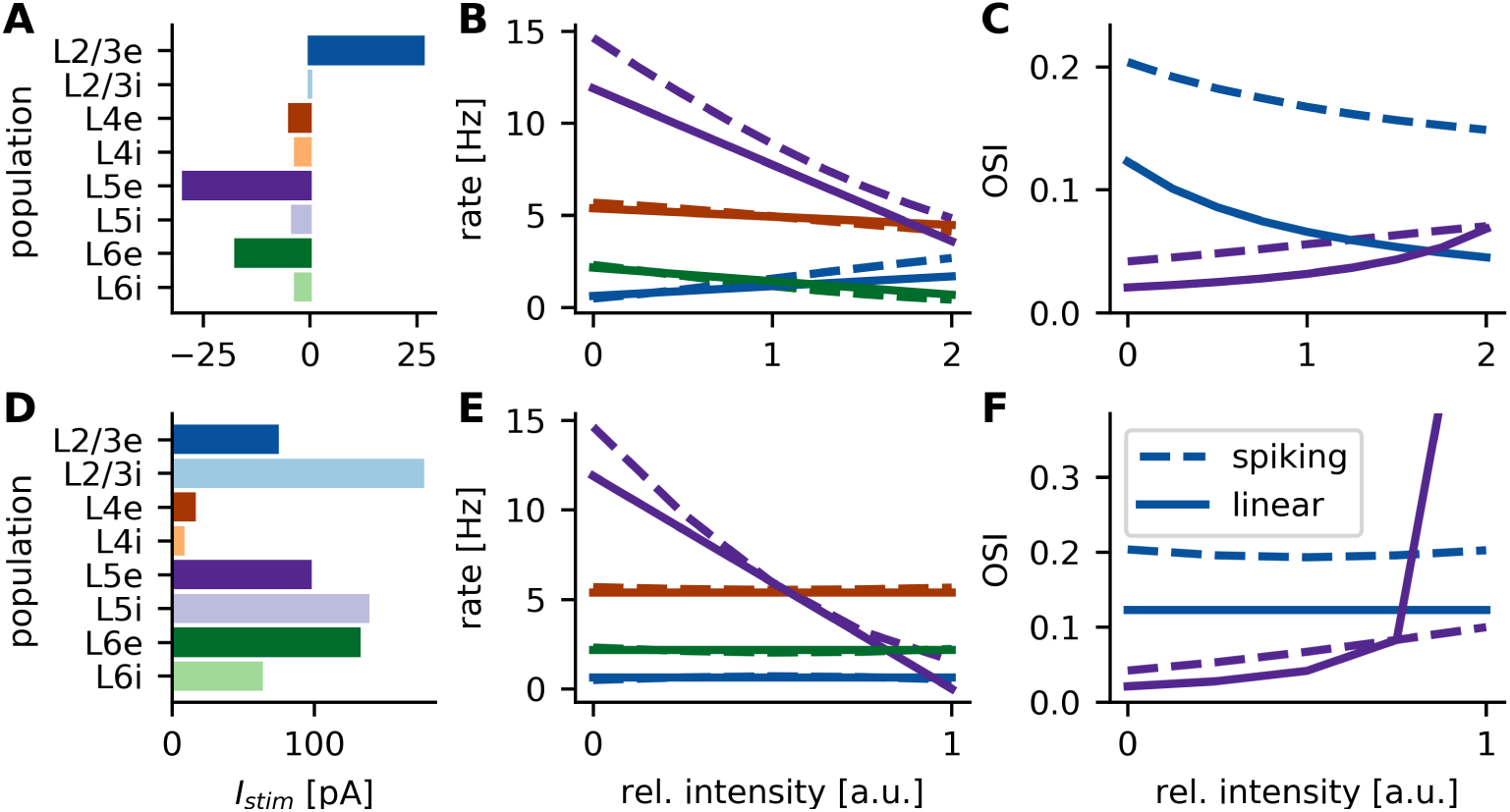
Predictions for future current stimulation experiments. **A-C** Current stimulus 1: Effective deletion of a distinct mode from the input. **D-F** Current stimulus 2: Selective suppression of L5e activity. **A**,**D** Single neuron current stimulation required to achieve the effect. **B**,**E** Change of the mean rate of excitatory populations upon current stimulation. Inhibitory populations are omitted for simplicity. x-axis shows current intensity relative to values in A,D, respectively. In all panels, lines represent the linear model (Model C), and dashed lines the spiking neural network simulations (Model A). **C**,**F** Change of orientation selectivity in L2/3e and L5e due to the two different current stimulations.

When the current is applied with an intensity that exactly removes the mode from the input, the change in output exactly matches the sum of the remaining modes (Fig 9B, relative intensity 1). For stronger stimulation intensities, the input mode is overcompensated. The mean population rates of the linearized network and spiking network model are in good agreement. Also in the spiking neural network model, the mean rates change almost linearly, even if the input mode is overcompensated by a factor of two.

As expected, the reduction of L5e firing rates leads to an increase in orientation selectivity of 250% from 0.02 to 0.07 in the linearized network model (75% in the spiking network model, Fig 9C). Simultaneously, due to the increase in firing rate in L2/3e, the orientation selectivity of this population decreases by 66% from 0.12 to 0.04 in the linearized network model (25% in the spiking network model). The discrepancy between the OSI of the spiking and linearized network model can be partially explained by the bias of random tuning due to the fluctuations of the spiking process, which were also the reason for non-zero OSIs observed in the spontaneous, unstimulated condition in Fig 4B. Other contributions could come from a weak crosstalk between the baseline and modulation pathways, or from deviations from the assumptions of Poisson firing statistics and negligible correlations.

Effectively deleting the mode from the input is not the only option to reduce the activity of L5e neurons. From our analytic calculations, all feed-forward and recurrent baseline gains are known. Exploiting this knowledge, we can design yet another current stimulus that exclusively affects L5e neurons, leaving the mean firing rates of all other populations unchanged (**Methods**). The desired change of rates for this stimulus can be conveniently summarized in a vector Δ*ν*^L5e^. This vector is zero for all populations except L5e. For this population, it contains the negative of the mean rates in the stimulation condition such that the resulting rates vanish. The required effective external perturbation can then be calculated by

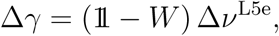

which are the external currents scaled with the known neuron sensitivities.

The required stimulation currents are shown in Fig 9D. Surprisingly, in this case, the input currents to L5e neurons all need to be positive, although the desired effect on the output rates is negative, and the rates of these neurons are reduced. It is also interesting that input currents to all neurons except L4 are of similar strength. It is clear that the other populations also require substantial stimulation to maintain their original rate, since the altered rate of L5e neurons needs to be compensated. However, the magnitude of these currents surprises as L5 does not serve as a major input to any other population within the network (Fig 1B, Fig 5A). This highlights again the strongly recurrent and often unintuitive nature of such networks.

Fig 9E shows that the new current stimulus successfully reduces the rate of L5e neurons, while keeping all other firing rates unchanged. Similar to the first current stimulus discussed above, the two models match quite well. Only for strong stimulation intensities, when the rates of L5e neurons approach zero, the two models show discrepancies. In this regime, the rates of L5e begin to be rectified. Since this is a non-linear effect, it can of course not be captured by the linearized version of the model.

Analogously to our first prediction for current stimulation, L5e neurons show a strong increase in orientation selectivity (Fig 9F). In the linear model, the OSI rises very sharply for large current intensities. This is an artifact of the neglected rectification of neuron rates, as already observed above. In the more realistic spiking network model (Model A), the OSI also increases strongly by about 150% (from 0.04 to 0.1). However, in contrast to the first current stimulus discussed above, in this case orientation selectivity of L2/3e neurons is not compromised by the additional input currents. The OSIs of the other excitatory and inhibitory populations also remain unchanged (data not shown).

## Discussion

### Model

The goal of this work was to study signal propagation across layers in primary visual cortex of rodents, with a particular focus on the emergence of orientation tuning in layers that have no direct access to thalamic input. We decided to employ a model that was developed previously by Potjans and Diesmann (2014). The synaptic connectivity defining this model was rigorously derived from the results of anatomical and electrophysiological measurements. The parameters describing neuronal connectivity depend on the pre-synaptic and post-synaptic population, but they do not depend on the functinal properties (tuning) of individual neurons. Furthermore, in order to focus our analysis on the impact of neuronal connectivity on dynamics, single-neuron parameters were chosen the same for all neurons, independently of the population they belong to.

Since not all required connectivity data are available, Potjans and Diesmann (2014) combined data from different species (cat and rat) into one single generic model circuit. In addition, the measurements were performed in several different brain regions, so it is debatable whether this model can account for information processing in mouse visual cortex at all. In view of this, it may be surprising that the predictions of our study with regard to neuronal activity are in very good agreement with direct electrophysiological measurements performed in mouse visual cortex. This finding is in fact compatible with the idea of a universal cortical circuit which is found in different species and in different cortical regions devoted to the processing of sensory information. It will be interesting to see, to which degree a neural network model based on the full mouse/rat visual cortex connectome (when it becomes available) at all deviates from the present model.

In our work, the model proposed by Potjans and Diesmann (2014) was extended by adding neuron-specific thalamic input, which is tuned to the orientation of a visual stimulus, i.e., a moving grating. This input was provided to both the excitatory and inhibitory populations in L4 and L6. From experiments, it is known that neurons modulate their firing rate with the temporal frequency of the grating (Niell and Stryker, 2008; Durand et al., 2016). However, in order to keep the already complex model and analysis as simple as possible, we did not include this aspect of visual processing into our considerations. In fact, our analysis showed that the cortical network operates in a quasi-linear regime. Therefore, we expect that the additional temporal modulation of firing rates would not have a significant impact on the mean values studied and reported here.

Via random connections, the very weak orientation bias in the input was enough to induce strong orientation tuning in L2/3, see Hansel and van Vreeswijk (2012) and Sadeh et al. (2014); Sadeh and Rotter (2014, 2015) for an analysis of this generic effect. Note, however, that we did not make any assumptions about the origin of the orientation bias in the input. It could be a consequence of the alignment of the receptive fields of afferent neurons (Hubel and Wiesel, 1962), or it could be inherited from already tuned thalamic neurons (Marshel et al., 2012; Piscopo et al., 2013; Scholl et al., 2013). While the input was assumed to convey information about a visual stimulus throughout the present study, our analysis of the layered network described here does not rely on this interpretation. Our approach could therefore also be applied to study feature selectivity in other sensory modalities, like in somatosensory or auditory cortex.

The distributions of orientation selectivity that emerge in our model across the different layers and neuronal populations are qualitatively very similar to experimental recordings (Heimel et al., 2005; Girman et al., 1999; Niell and Stryker, 2008; Sun et al., 2016; Durand et al., 2016) (see also (Van den Bergh et al., 2010) for some inconclusive results). This represents a remarkable result, as the model is based on measured connectivity in different cortical regions in different species, and it was not at all designed and tuned as a specific model of rodent visual cortex. Furthermore, the recordings cited above were performed in adult mice, which are known to have some degree of feature-specific connectivity. Our model, in contrast, entirely lacks specific connectivity, resembling the situation at eye opening (Ko et al., 2013). Our analysis demonstrates that well-tuned neuronal responses can emerge in all cortical layers without feature-specific connections between neurons. On the other hand, it is known that specific connectivity can work as a concurrent mechanism to improve orientation selectivity and yield the higher values observed experimentally (see also (Sadeh et al., 2015)).

While the distribution of firing rates in spontaneous and evoked conditions are qualitatively similar to experimental findings (Niell and Stryker, 2008; Durand et al., 2016), there are also discrepancies in several respects (Fig 2D/E). It is difficult to asses the impact of these differences for the findings of our work. However, our results are generally robust to changes in central network parameters, despite considerable rate changes. Therefore, we expect that the neuronal mechanisms described in our work still apply when more precise and detailed models of the microcircuit are developed.

In the present model, the mouse which is subject to visual stimulation is assumed to be non-moving. As shown in a recent study (Durand et al., 2016), neuronal responses in primary visual cortex of rodents are very similar in awake and anesthetized animals. In contrast, the responses dramatically change during locomotion behavior (Niell and Stryker, 2010). Different inhibitory neuronal subtypes play a central role in this (Pakan et al., 2016). A future enhanced version of our model might account for such effects as well.

### Microcircuit perspective

While analyzing network activity in terms of firing rates and orientation tuning, it becomes apparent that a direct interpretation of connectivity cannot fully explain all features of the dynamics observed in network simulations. For example, although afferent connectivity originating from L4e is very similar, firing rates and orientation selectivity in L2/3e and L5e are quite different. As the input is comprised of both excitatory and inhibitory projections that partially cancel each other, a small difference in the input may have strong effects on the operating point the two populations eventually settle in (Van Vreeswijk and Sompolinsky, 1996, 1998). Such effects are not straight-forward to predict, since in a highly recurrent network neuronal output simultaneously also provides input to the same network. Sometimes this leads to non-intuitive effects like effective inhibitory influence of excitatory neuron populations (Tsodyks et al., 1997). To master such difficulties in the analysis, a system-level view of complex micro-circuits is inevitable (Yuste, 2015; Bassett and Sporns, 2017).

### Baseline and modulation pathways

Previously, inhibition dominated recurrent networks were shown to exhibit two distinct processing pathways (Sadeh et al., 2014). While the untuned and uninformative component (baseline) of the input is strongly attenuated by the recurrent network, the modulated component, which conveys all the information about the stimulus, has a much higher gain. This results in a net amplification of feature selectivity through the input-output transformation exerted by the network.

By separating the (linearized) layered model into two systems, we were able to analytically calculate the feed-forward and recurrent gains of the baseline pathway also for a multi-population system. In combination, these two gains describe how the input to one population affects the output of any other population. In contrast to our model, neurons in the model of Potjans and Diesmann (2014) have binomially distributed in-degrees, leading to a certain degree of cross-talk between the two pathways by non-vanishing interference terms in Eq. 12. While this introduces some discrepancies between the direct and separate solutions of the linear model, it does not impair the described mechanism.

It should be emphasized that the separation between baseline and modulation pathways, its different gains, and thus also tuning amplification are purely linear effects. This is possible, as a linear transformation may impose different scaling factors to different input components.

### Mode decomposition

Instead of analyzing the operation of a network from the view-point of neuronal populations, we chose a basis that is more natural for the network in question. We decomposed the input into eigenmodes of the system, where the eigenvalues of the effective connectivity correspond to the gains of the individual input modes. Interestingly, the high gain of L5e can be attributed to a single mode, which has a much higher gain than all the other modes. Due to this mode, L5e neurons exhibit a high firing rate and a weak selectivity for stimulus orientation.

What is the functional significance of having a mode with this exceptionally high gain? For a preliminary answer, it is important to take into consideration that the different populations in the layered cortical network play different roles for information processing. For instance, L2/3e neurons project to higher brain areas and need to transmit information in a reliable way (Felleman and Van, 1991). Therefore, it is desirable that this population has a strong attenuation of the baseline and, thus, strong tuning amplification. L5e neurons, on the other hand, mainly project to sub-cortical brain regions, including thalamus (Sherman, 2016). It is likely that this population is involved in feedback control mechanisms, where tuning information may be less relevant.

### Predictions for optogenetic intervention

Using the theory developed in this work, we are in the position to devise external (e.g. optogenetically applied) stimuli which effectively manipulate the orientation tuning of L5e neurons. We designed two different such stimuli which reduce the mean firing rates of these neurons, thus increasing their tuning. Interestingly, although the effect on the rate of L5e is identical, one stimulus requires a positive current, while the other utilizes a negative current. This can be explained by the indirect effect of the stimulus on other populations, propagated via the recurrent network. Our calculations constitute strong predictions of our model, which can be tested with some effort in experiments. Different subpopulations can be stimulated by a combination of channelrhodopsin and halorhodopsin with different wavelength sensitivities, as described by Klapoetke et al. (2014), possibly in combination with locally confined optical stimulation.

Throughout our study, and in particular for the derived predictions, the consistency of results between the spiking network and the linear model are remarkable. Only for very strong stimulation, when non-linear rectification sets in, some deviations become evident. This confirms previous findings that a network of highly nonlinear neurons can be effectively linearized about its dynamical operating point, which corresponds to the balanced state of the excitatory-inhibitory network (Van Vreeswijk and Sompolinsky, 1996, 1998).

In our work, the newly developed mathematical tools were applied to design current stimuli which effectively and selectively suppress L5e activity. In our model, all rate gains are known from the decomposition into baseline and modulation pathways. Therefore, other types of experiments can be performed as well, even for complex multi-population networks different from the primary visual cortex. This makes the suggested method an ideal tool for the design of external current stimuli in a large variety of experiments.

In summary, we presented a novel method for analyzing the dynamics of multipopulation networks in terms of their input-output relations. We applied the approach to the analysis of a layered model of rodent visual cortex, revealing interesting and to some degree also unexpected and non-intuitive consequences of the underlying connectivity. In addition, we derived predictions for future optogenetic experiments, which could be performed to test the power of our computational analysis.

## Acknowledgments

This work was partially supported by the European Union’s Seventh Framework Programme (FP7/2007-2013) under Grant Agreement 600925 (NeuroSeeker), by the BMBF (grant BFNT 01GQ0830) and DFG (grant EXC 1086). The HPC facilities are funded by the state of Baden-Württemberg through bwHPC and DFG grant INST 39/963-1 FUGG. In addition, our work was promoted by the German Academic Exchange Service (DAAD) and the Carl Zeiss Stiftung. We thank Uwe Grauer from the Bernstein Center Freiburg as well as Bernd Wiebelt and Michael Janczyk from the Freiburg University Computing Center for their assistance with HPC applications.

## Code Accessibility

Simulation and analysis code is available upon request.

## Conflict of interests

The authors declare no competing financial interests.

## Supporting information

**sFig 1:**
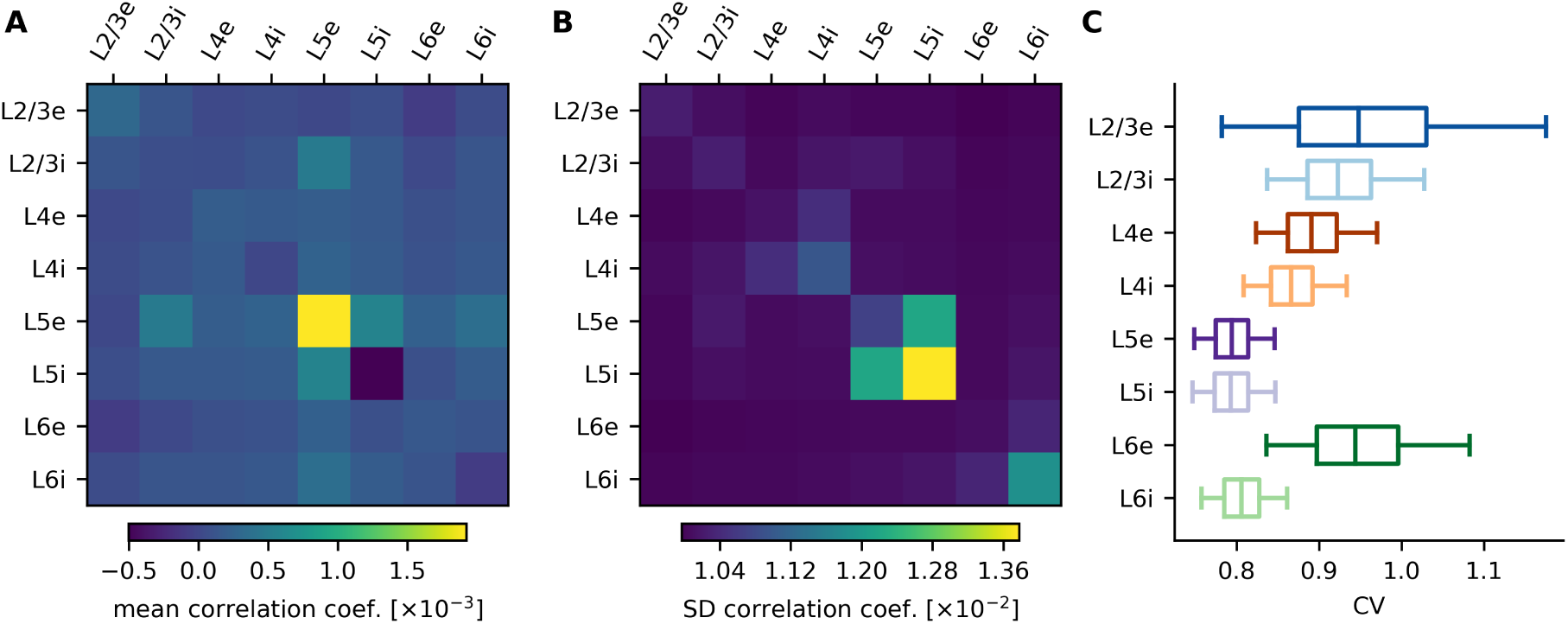
Correlation and spike regularity in the spontaneous condition. **A-B** Mean and standard deviation of spike count correlations between neurons from different populations. A bin size of 10ms was used. **C** Distributions of coefficient of variation of inter-spike-intervals for neurons in different populations. Same quantiles as described in Fig 2 are shown.

**sFig 2:**
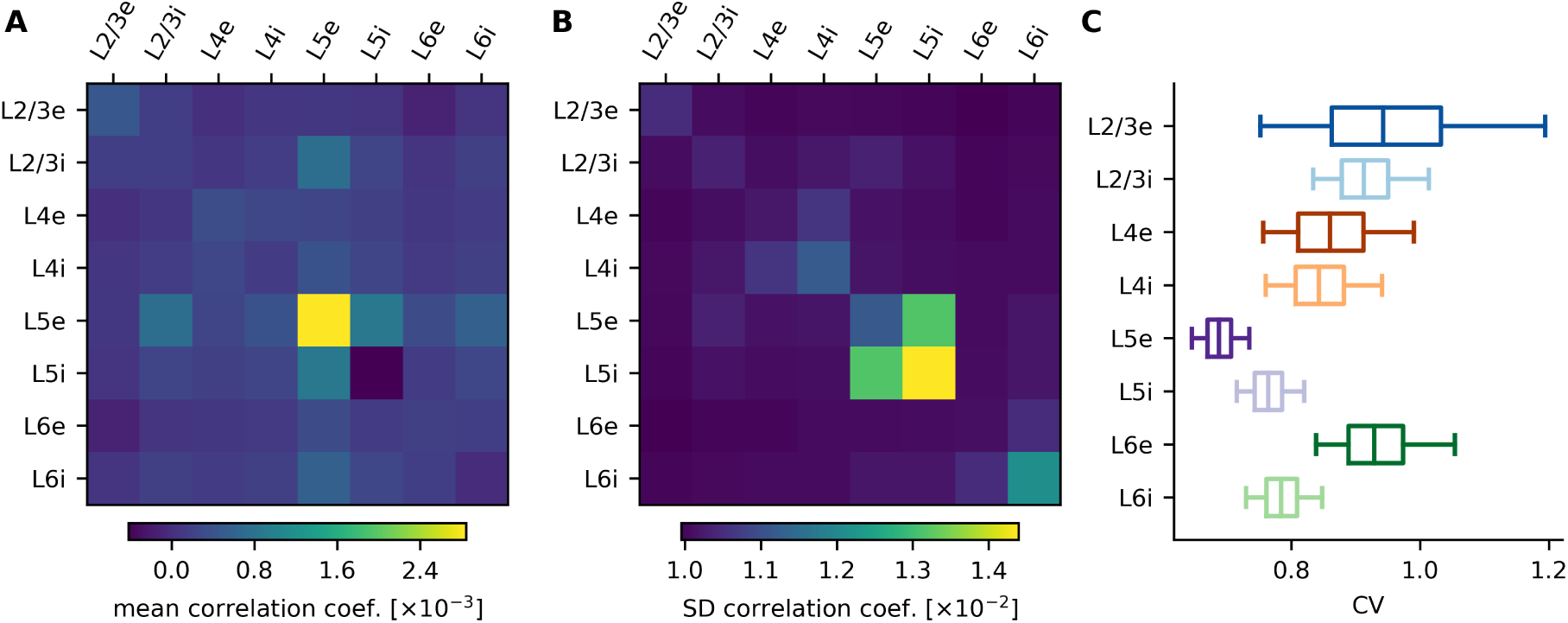
Correlation and spike regularity in the stimulated condition. Same values as in SFig 1 are shown for the stimulated condition.

**sFig 3:**
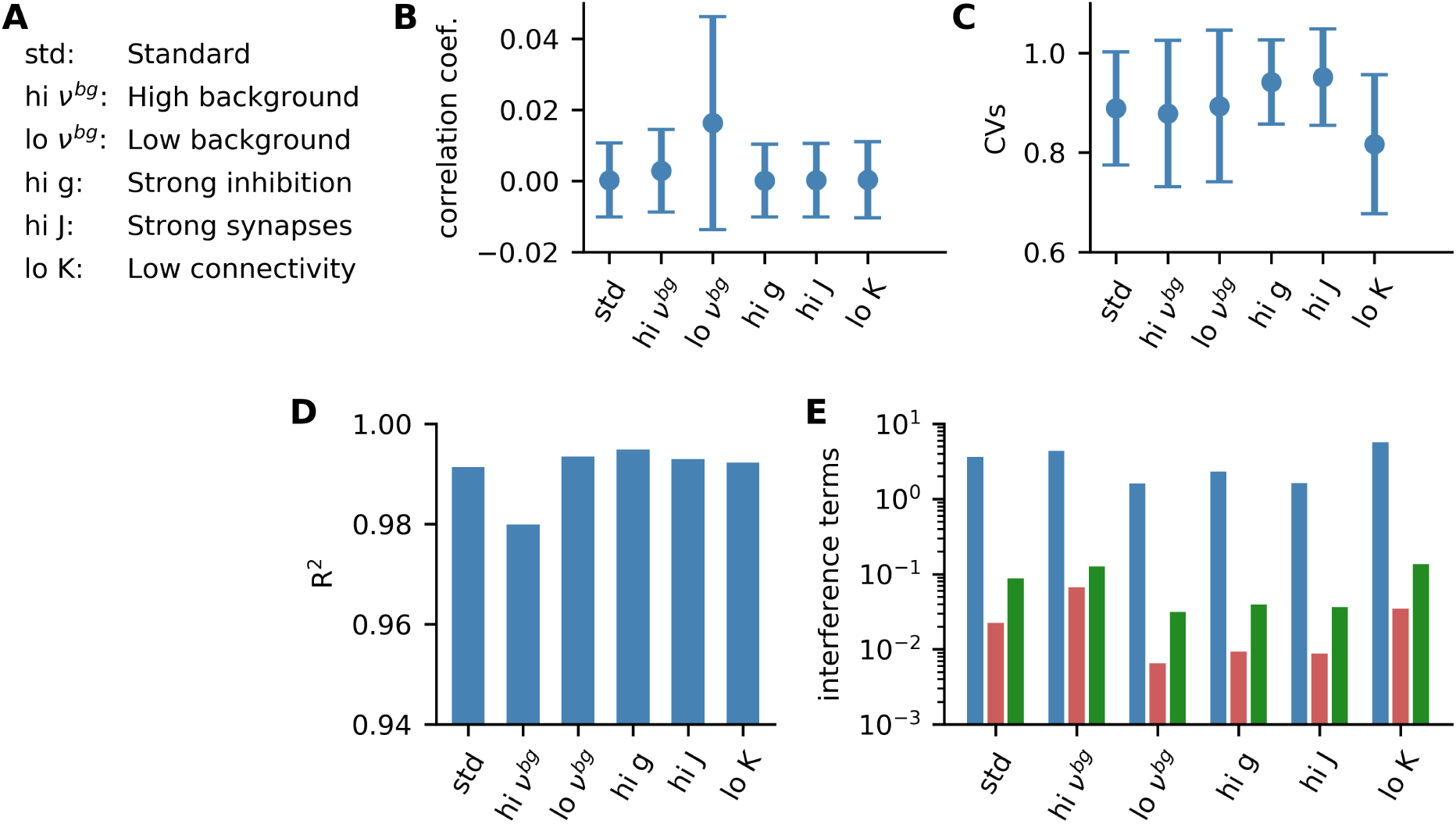
Characteristic quantities compared over parameter sets. **A** Abbreviations used in other panels. **B** Mean and SD of correlation coefficient over all populations. **C** Mean and SD of CV of inter-spike-intervals over all populations. **D** Coefficient of determination (*R*^2^) quantifying fit between direct linear solution and baseline and modulation decomposition. **E** Magnitudes of interference terms in Eq. 12 on logarithmic scale (blue: Δ*β*, red: *Q*Δ*ν*_*M*_, green: *S*Δ*ν*_*B*_).

**sFig 4:**
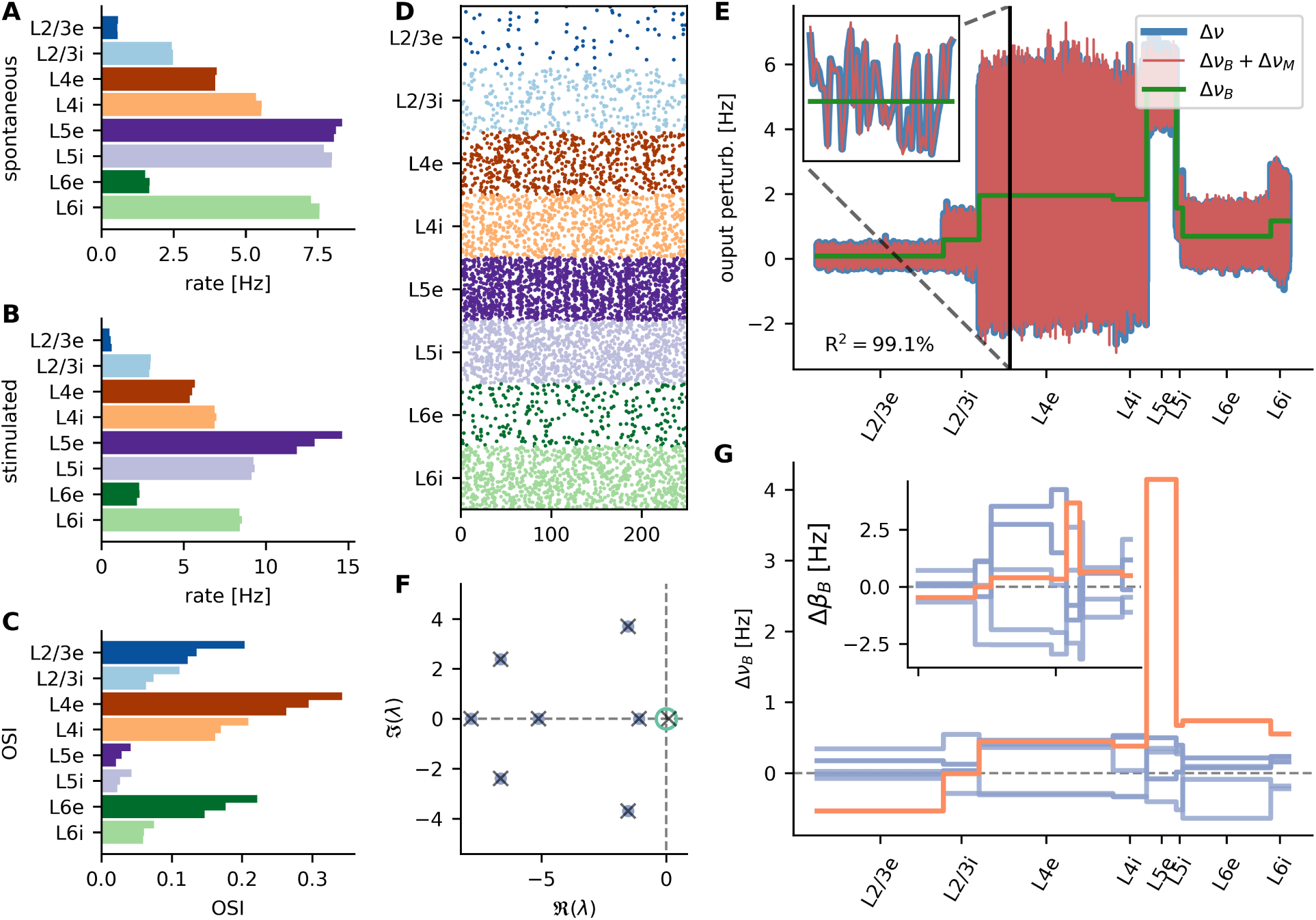
Central results summarized for standard parameter set. **A/B** Mean firing rates for spontaneous and stimulated condition. For each population, the results of the spiking model (Model A, top), rate model (Model B, middle) and linear model (Model C, bottom) are shown. **C** Mean orientation selectivities for stimulated condition from all three models. **D** Raster plot for the stimulated condition for 250ms of 500 neurons in each population. **E** Comparison of baseline and modulation solution with direct solution of the linear model (c.f. Fig 6C). **F** Eigenvalue spectrum of the effective connectivity *W* (c.f. Fig 7B). **G** Input perturbation Δ*β*_*B*_ (inset) and output perturbation Δ*ν*_*B*_ decomposed into input and output modes, respectively (c.f. Fig 7D).

**sFig 5:**
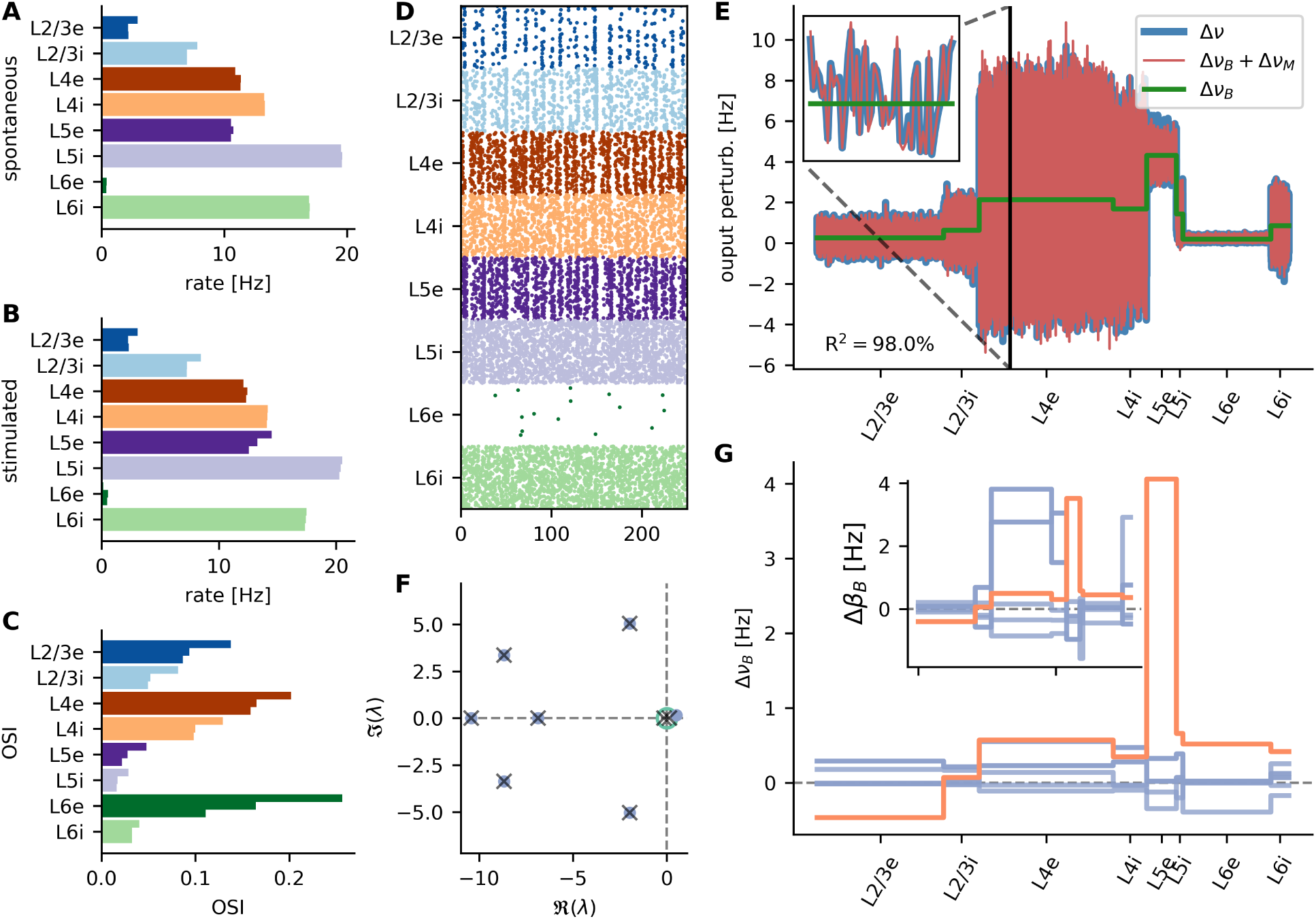
Central results summarized for high background parameter set. See SFig 4 for a description of the individual panels.

**sFig 6:**
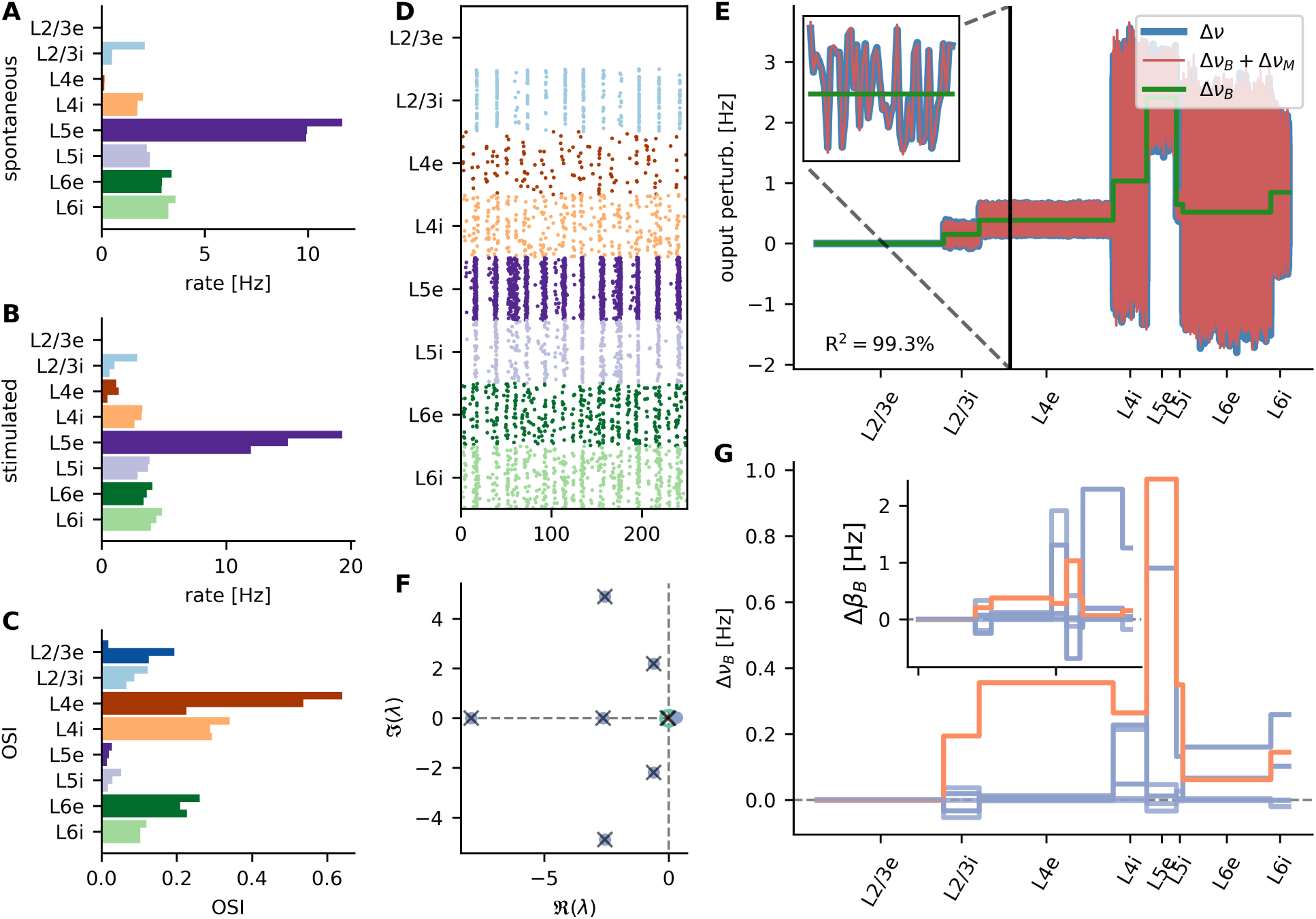
Central results summarized for low background parameter set. See SFig 4 for a description of the individual panels.

**sFig 7:**
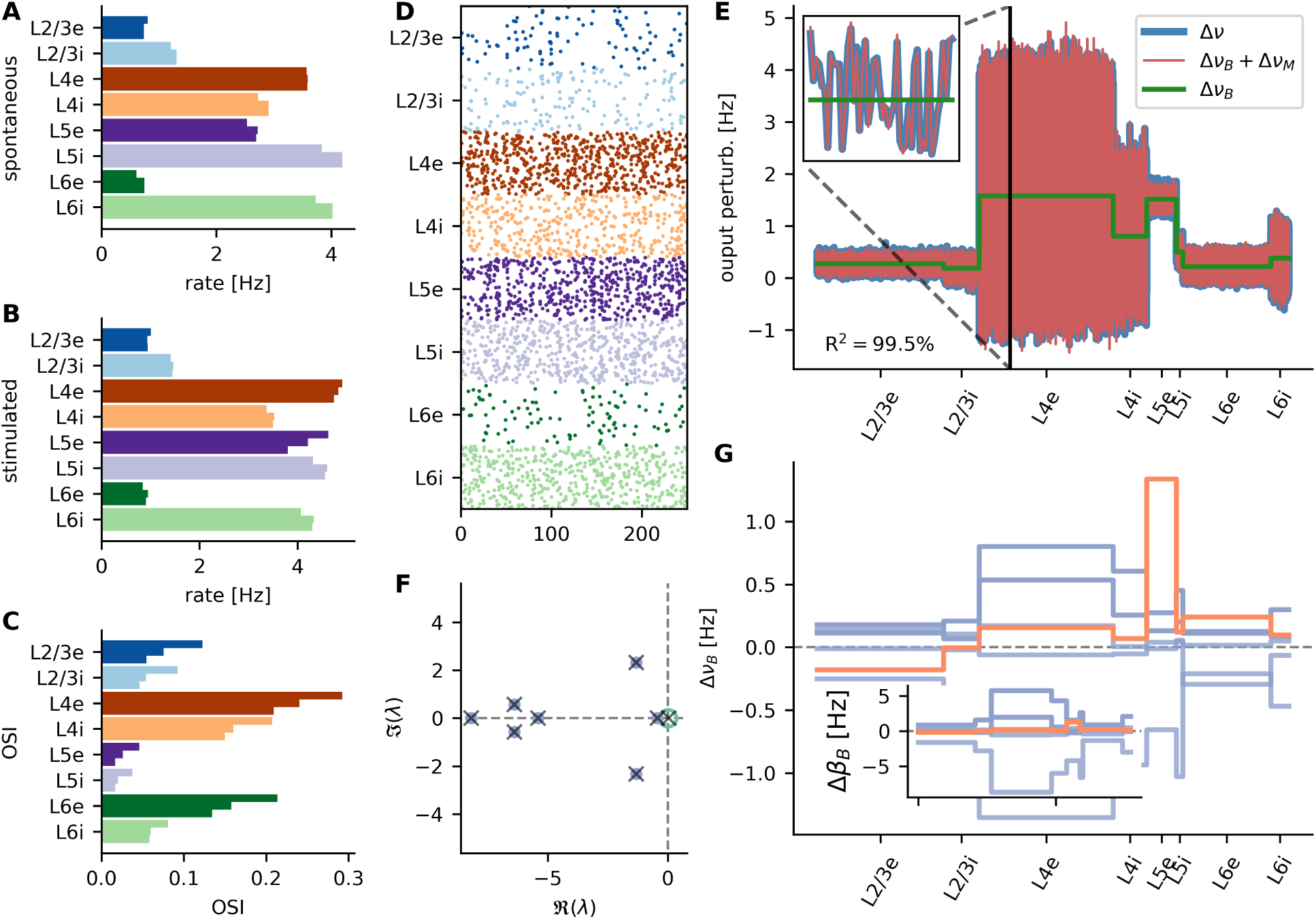
Central results summarized for strong inhibition parameter set. See SFig 4 for a description of the individual panels.

**sFig 8:**
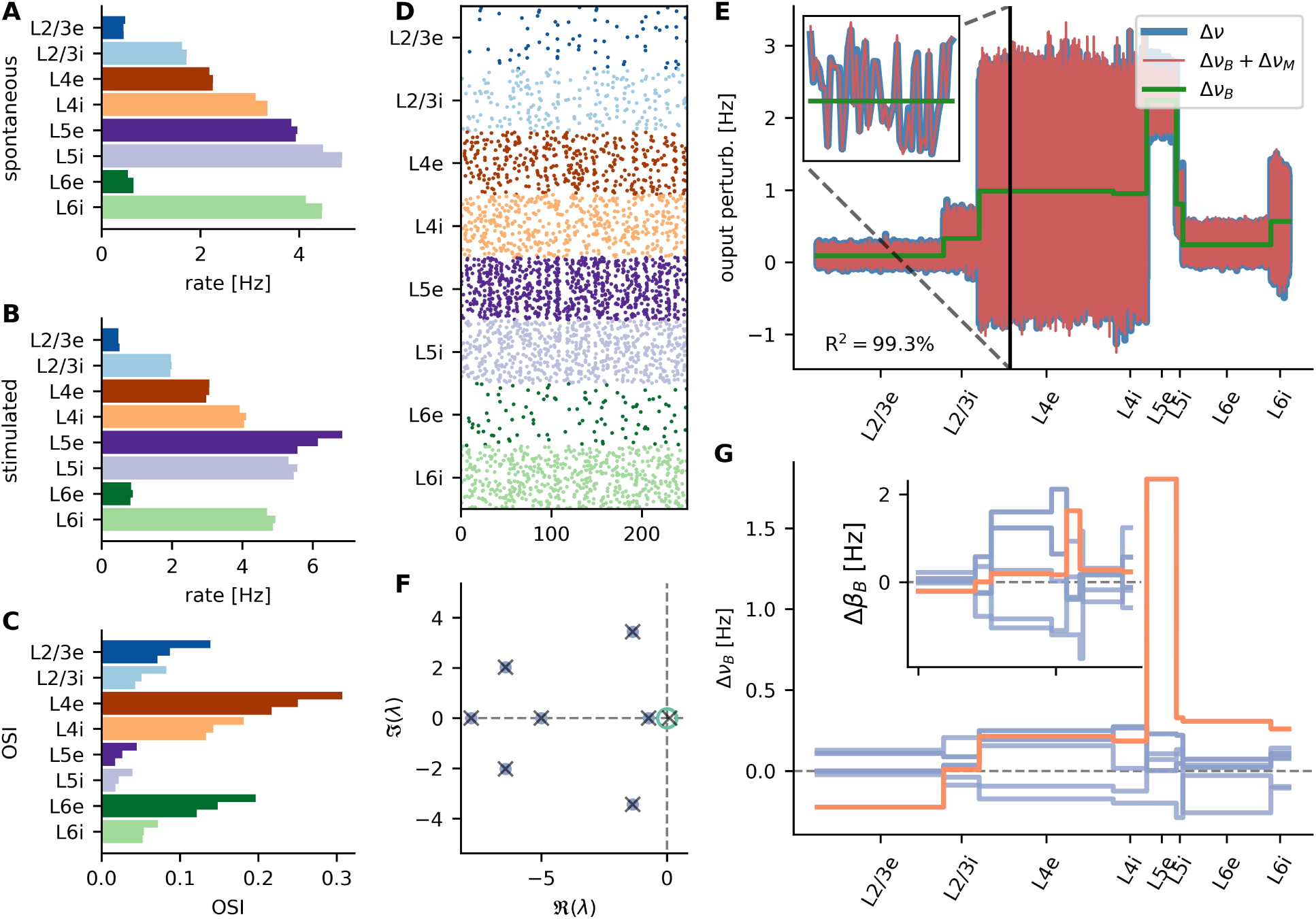
Central results summarized for strong synapses parameter set. See SFig 4 for a description of the individual panels.

**sFig 9:**
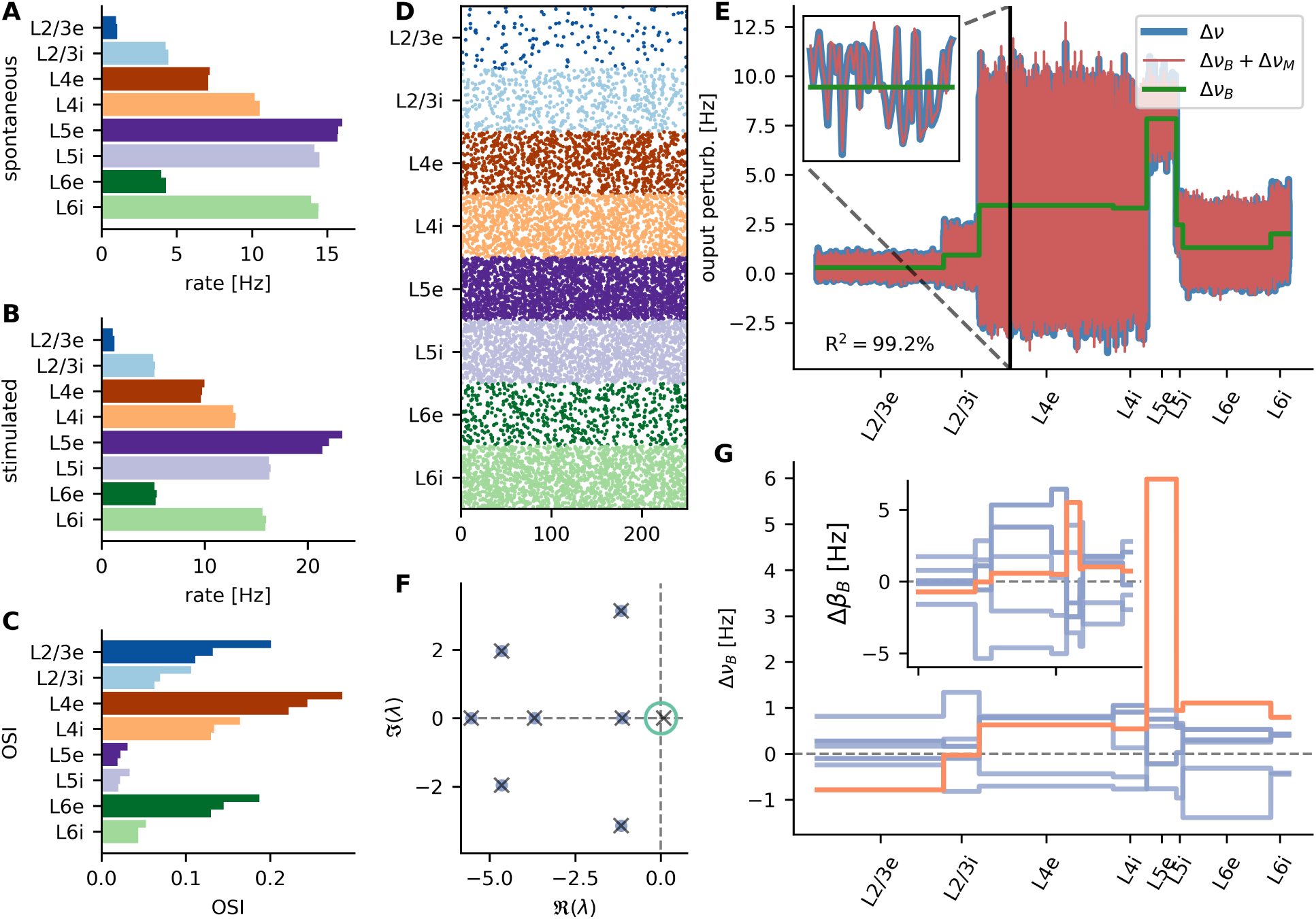
Central results summarized for low connectivity parameter set. See SFig 4 for a description of the individual panels.

**sFig 10:**
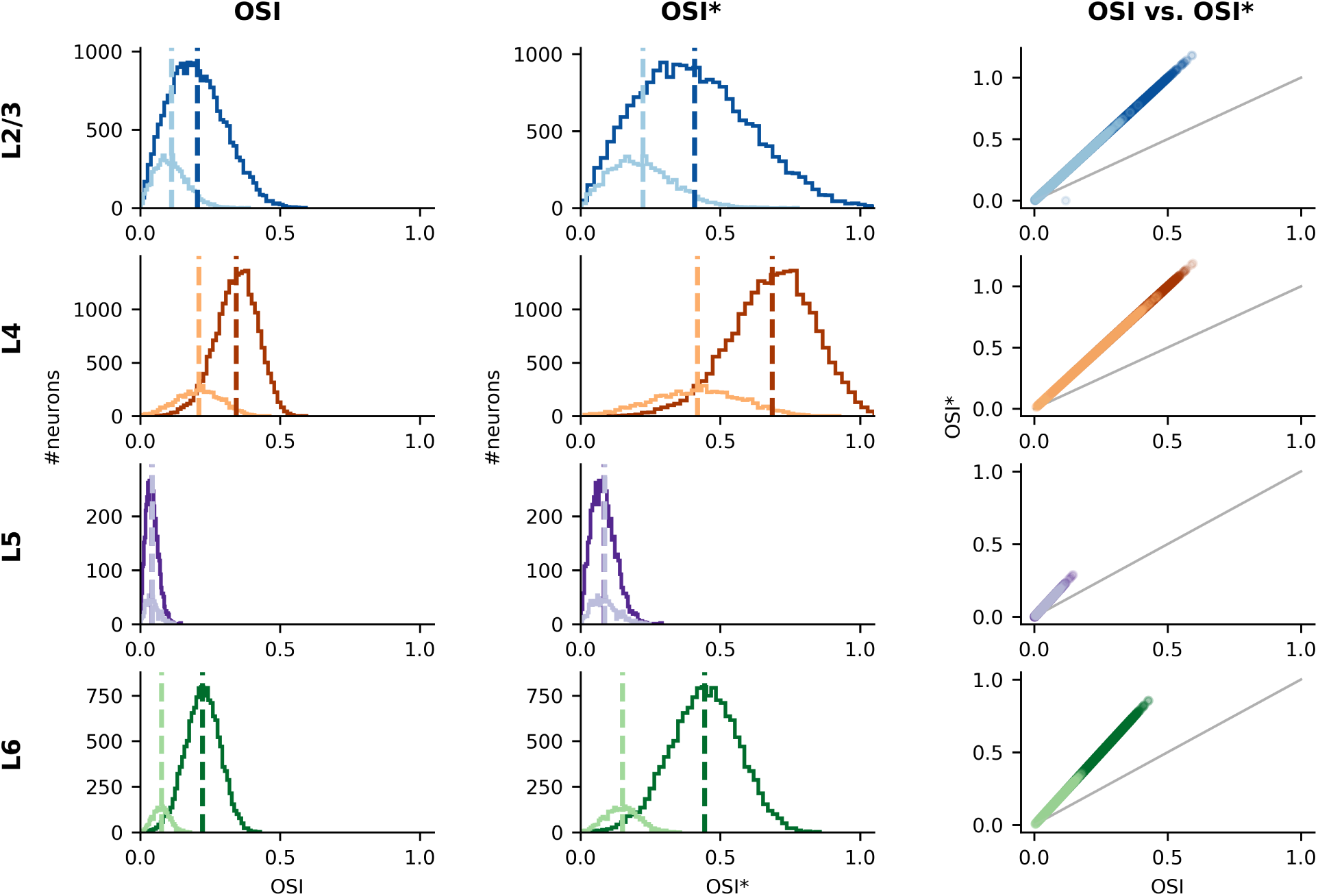
Comparison of orientation selectivity measures. Left column: Distribution of orientation selectivity measure based on circular statistics (OSI). Middle column: Orientation selectivity measure based on cosine fit (OSI*). Right column: Comparison of the two orientation selectivity measures. Each dot represents a single neuron in the respective population.

**sFig 11:**
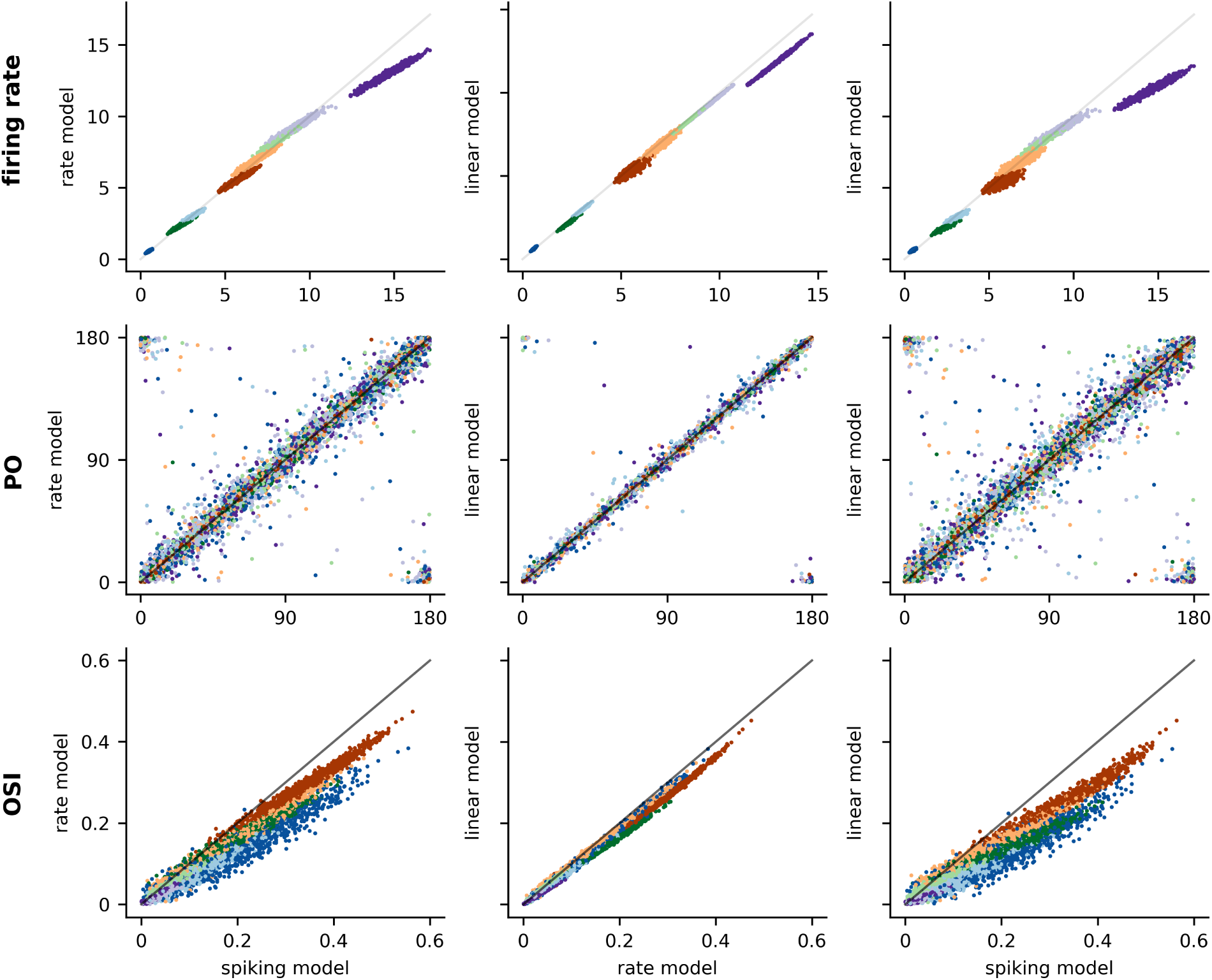
Model comparison. Comparison of the three central observables of the different network models. Top row: single neuron firing rates, middle row: preferred orientation (PO), bottom row: orientation selectivity index (OSI). The first column compares the spiking network model (Model A) with the firing rate model (Model B), the second column compares the firing rate model with linearized network model (Model C) and the third column compares the spiking network model with the linearized network model. Same color code is used as in previous figures to indicate the eight different populations.

